# Episodic memory encoding by a place cell sub-population

**DOI:** 10.1101/682831

**Authors:** Jake Ormond, Simon A. Serka, Joshua P. Johansen

## Abstract

Study of the hippocampal place cell system has greatly enhanced our understanding of memory encoding for distinct places, but how episodic memories for distinct experiences occurring within familiar environments are encoded is not clear. One possibility is that different place cell populations encode details of the novel experience or maintain the representation of the unchanged environment. We developed an aversive spatial decision making task which induced partial remapping in CA1, allowing us to identify both remapping and stable cell populations. We found that remapping cells exhibited distinct features not present in stable cells. During memory encoding, their theta phase preferences shifted to earlier phases, when CA3 inputs are strongest. Further, their recruitment into replay events increased during learning, unlike that of stable cells. Our demonstration of a sub-population of place cells identified on the basis of their degree of remapping and exhibiting unique changes in their spike firing properties with learning lend support to a model in which novel and familiar spatial/contextual information is encoded and maintained, respectively, by separate place cell populations.

## INTRODUCTION

Episodic memories include details about salient experiences from our past as well as information about the location in which these events occurred^1^. The hippocampus plays a central role in episodic memory and spatial decision making^2–5^, and this is thought to rely in part on its ability to create and store unique representations or maps of different spatial and non-spatial contexts^6, 7^. Alteration of the spatial representation, termed remapping^8^, is also thought to play a role in the encoding of episodic events occurring in familiar environments^9^. How this occurs without disrupting the representation of the physical and spatial features of the environment is not clear. Furthermore, the mechanisms through which episodic experiences are integrated into hippocampal representations during learning are not well understood.

Studies in which physical properties of the environment, such as color or shape of the recording apparatus, are altered have shown that the infield firing rates of existing place fields can change in response to changing sensory input without any change in place field location, a process termed rate remapping^10^. Further support for this concept has been found in studies in which changes in behavioral contingencies, such as turning direction on a plus or T-maze, also produce rate remapping^11–13^. However, studies of remapping during episodic memory encoding using reward or fear learning have reported changes in the place field locations of some, but not all, the recorded place cells ^14–18^.

Partial remapping, meaning remapping in a subset of the place cell population, suggests the possibility of a unique population of place cells which might dynamically encode information relating to events, while leaving a more spatially specific population to stably encode space. But how might episodic memory encoding take place such that it is restricted to one subpopulation, while leaving another unchanged? A widely held theory of hippocampal encoding and recall for spatial learning in CA1 posits that encoding is driven by entorhinal inputs conveying information about the real-time state of the external world, while recall is driven by CA3 inputs relaying information stored within the CA3’s recurrent network^19, 20^. Because entorhinal inputs are strongest on the ascending wave and peak of theta, whereas CA3 inputs are strongest on the descending wave^21–23^, this theory further posits that encoding and recall will be similarly segregated with respect to theta. Thus, one possibility is that the efficacy of entorhinal inputs to the encoding subpopulation may be selectively increased, and this may be visible as a shift in the spike times of the member cells towards the ascending wave and peak of theta. Another possibility is that encoding cells may be preferentially recruited into sharp wave ripple (SWR) associated replay events. Recent work has shown that in CA1, a distinct population of place cells can be identified on the basis of differential recruitment into replay events^24^. Further, patterns of activity associated with spatial novelty or reward learning have been shown to recur more frequently in these events^25, 26^. Last, these events have been shown to be important for the stabilization of place fields^27^, and for the consolidation of place cell assemblies in novel environments ^28^. However, it is not clear from this previous work whether these memory encoding and consolidation mechanisms occur in a specific population of hippocampal neurons.

We set out to explore the existence of an episodic encoding sub-population of place cells in CA1 and to determine the neural coding mechanisms through which these cells may be selectively integrated into hippocampal networks during learning and memory consolidation. Using a novel spatial decision making task incorporating avoidance of aversive stimuli, we found that remapping cells had unique phase locking properties, even before contextual learning, suggesting they were pre-selected for memory encoding. Their theta phase preference, rather unexpectedly, was shifted towards the descending wave, when CA3 inputs were strongest, and this shift increased during memory encoding. This shift was strongest during episodes of elevated power in the slow gamma band, further implicating CA3 inputs^21, 23^. Lastly, we found that during learning, remapping cells, but not stable cells, increased their ripple-centered firing rates and their participation in awake replay events.

## RESULTS

In reward learning paradigms, remapping occurs mainly around reward locations ^15, 29, 30^. Thus, any effort to determine what intrinsic properties cause some place cells to remap while others do not, is bound to be confounded by the locations of the cells’ place fields. To circumvent this limitation, we developed a novel decision making task incorporating aversive stimuli. Aversive stimuli produce spatially distributed remapping^17, 18, 31^, and require animals to use spatial knowledge about a stable, familiar environment in conjunction with information about changing episodic experiences occurring within this environment, leading to the creation of powerful episodic memories. Rats were trained to run laps on a track on which they could freely choose, or alternatively be forced, to run along 1 of 3 choice arms to return to a goal location for food reward (Fig. 1a-c). Rats (n = 4) were implanted with eye-lid shock wires, and shock (a 1 sec train of 1 msec pulses at 7 Hz, with intensity between 0.4 – 1.5 mA) could be triggered by breaking an infrared beam on the choice arms (Fig. 1d). During initial training, rats learned to navigate to the reward location; animals were never exposed to shock during this initial training. Once dCA1 tetrodes reached their target (962 place cells from 4 animals, 29 – 96 place cells per session; Fig. S1a), we ran a series of four sessions. Each session began with a baseline period using the previous session’s behavioral contingencies, which, in the learning sessions, was followed by a change in the contingencies, forcing the animal to adapt its behavior to avoid shock delivery; in control sessions, contingencies remained unchanged after the baseline (Fig. 1e, Fig. 2a). The session types were (in order; Fig. 1f): 1) noShockCTRL, in which no arms were paired with eye-lid shock; 2) Learning1, in which shock was introduced onto two of the choice arms; 3) shockCTRL, in which the contingencies from the Learning1 session were maintained; and 4) Learning2, in which the identities of the previous safe arm and one of the shock arms were switched (Fig. 1f; Fig. 2a). All animals learned the new task contingencies, showing a clear preference for the “safe” arm after an initial variable number of error trials (Fig. 1g-i; note that performance was initially below chance because the animal’s least preferred arm, determined from free choice trials at the beginning of the session, was always used as the subsequent safe arm; individual session learning curves in Fig. S1b). Further, rats showed long-term memory for the task contingencies, as demonstrated through a clear preference for the previous session’s “safe” arm at the outset of each new session (8-24 hours after the previous session; Fig. 1h, i).

**Figure 1.**
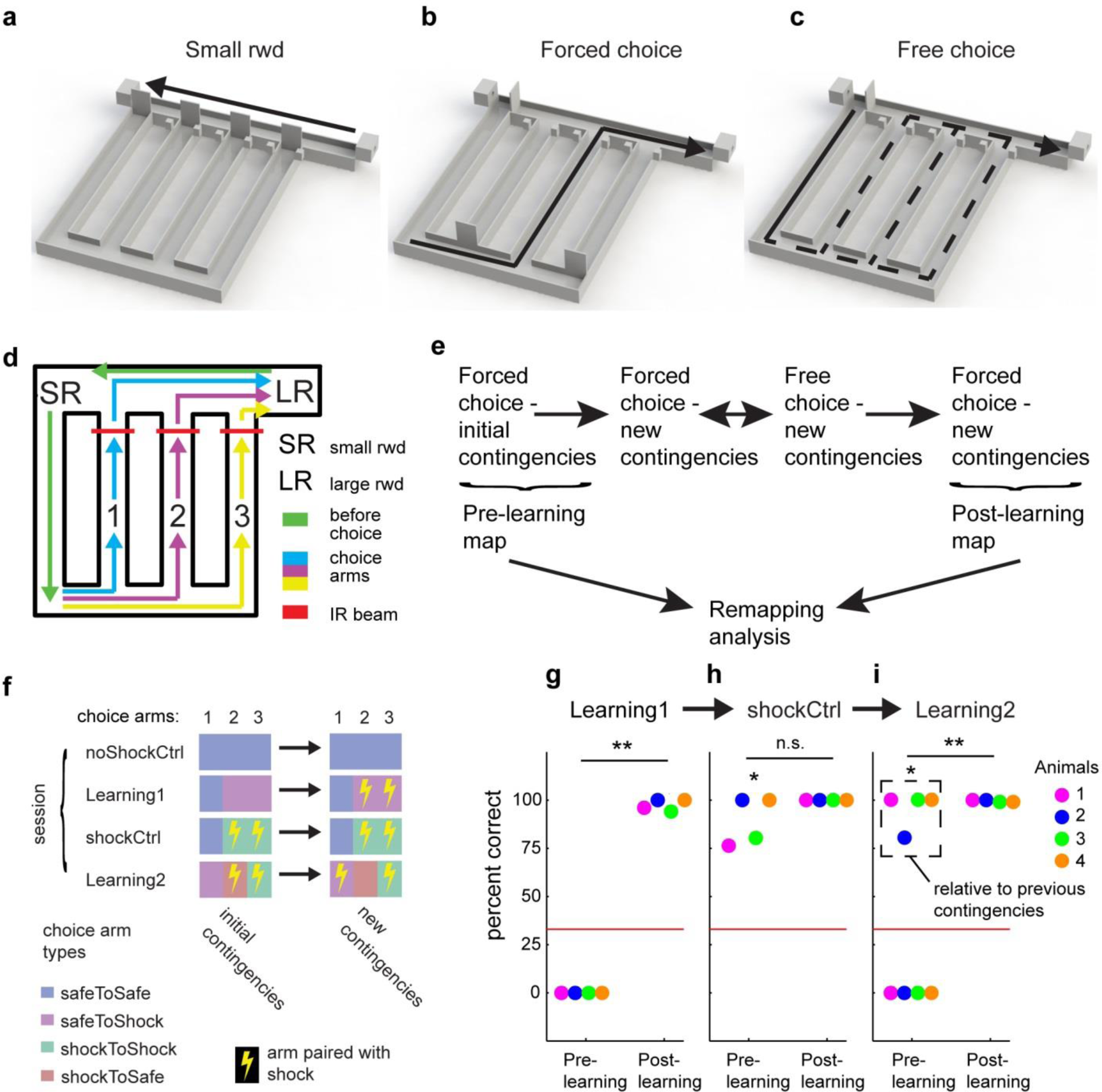
A novel aversive spatial decision making with aversive stimuli. (**a-c**) On each trial, animals first run to a small reward location (**a**) and then, from an initial start location at lower left corner (**b**), return to the main large reward location along 1 of the 3 choice arms. Sampling of an arm could be forced by closing off the other arms (**b**), or could be free choice (**c**), allowing an assessment of the animal’s arm preference. (**d**) For analysis, the maze was divided into 4 equal length segments (each segment is uniquely color coded) consisting of the path leading up to the choice points (“before choice”) and the 3 choice arms. Eye-lid shock was triggered by IR-beam breaks on the central arms. (**e**) The sequence of trial types within a session, and their use for the remapping analysis. Note that in a subset of sessions where the animal did not appear to switch its behavior after the first set of “Forced-choice-new contingencies” trials, a second set was then run, hence the recursive arrows (see Supplementary Fig. 1). (**f**) The four different session types with initial and final contingency combinations. (**g**) In the Learning1 sessions, animals (4 rats, 1 session each) consistently learned to avoid the shock arms in favor of the safe arm (*t*(3) = −65.7, *p* < 0.001). (**h**) Animal’s (4 rats, 1 session each) consistently remembered the Learning1 safe arm in the subsequent shockCtrl session (*t*(3) = 8.9, *p* = 0.003). No additional learning was observed across shockCtrl sessions (*t*(3) = −1.7, *p* = 0.1844). (**i**) During the Learning2 sessions, animals (4 rats, 1 session each) initially remembered the previous contingencies (dashed box; *t*(3) = 12.6, *p* = 0.0011), and then subsequently learned to take the new safe arm (*t*(3) = −344.7, *p* < 0.001). Note that performance in safeToShock and shockToNewShock sessions was initially below the chance level of 33% correct because the animal’s least preferred arm, determined from free choice trials at the beginning of the session, was always used as the subsequent safe arm.

**Figure 2.**
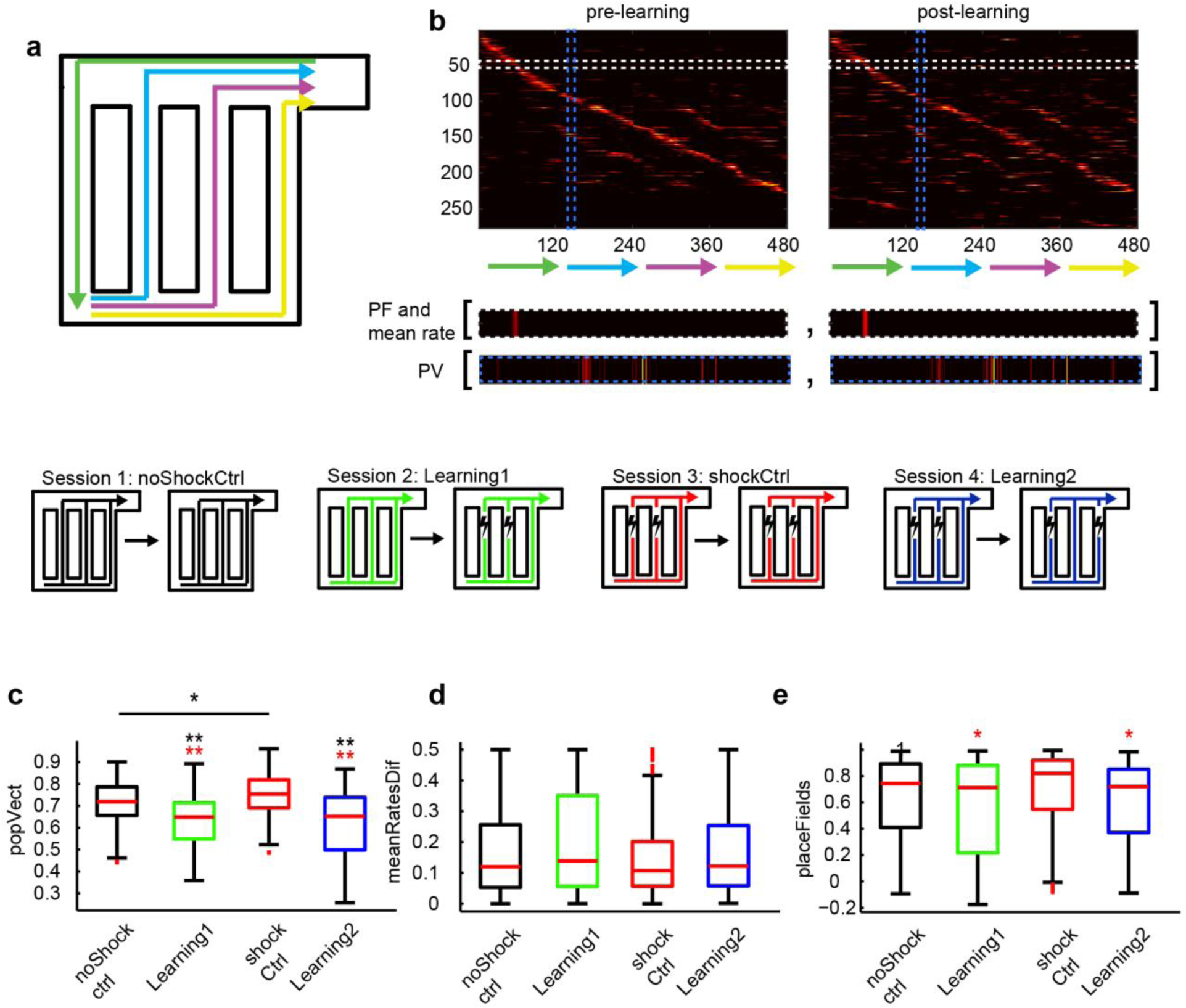
Remapping during task learning. (**a**) The 4 path segments run by the animal. (**b**) These 4 segmenets are linearized during analysis for the purpose of creating rate maps for all cells (top). To calculate the degree of remapping, correlation coefficients are calculated for pairs of rate maps (for individual cells; in white dashed boxes) or population vectors (representing the firing of all cells at a single spatial bin; purple dashed line) taken from before and after learning; additionally, changes in mean firing rate are also calculated for individual cells. The displayed rate maps are taken from the 272 cells recorded during the SafeToShock sessions which were classified as place cells. (**c**) Box plots of population vector correlation data. Kruskal Wallis test, *H*(3) = 147.9, *p* < 0.001, *n* = 200 spatial bins per session type. (**d**) Mean rate change analysis. *H*(3) = 6.6, *p* = 0.086, *n* = 228 (noShockCtrl), 277 (safeToShock), 218 (shockCtrl), 239 (shockToShock). (**e**) Place field correation analysis. *H*(3) = 16.6, *p* < 0.001, *n* as in (**d**). * or ** directly above box indicates significant difference from noShockCtrl (black), shockCtrl (red). Horizontal lines in (**c**) indicate significant difference between noShockCTRL and shockCTRL.

### Aversive experiences produce population level remapping

To characterize remapping resulting from task learning, we linearized the baseline and post-learning rate maps (Fig. 2a, b), and compared them using three common measures of stability (Fig. 2b): 1) correlations of population vectors (PV) representing the firing of all place fields at each spatial bin, 2) correlations of individual place cell rate maps (place field correlation), and 3) changes in individual place cell mean firing rates. The extent of remapping was assessed by comparing decorrelation and mean rate change in the learning sessions to the control sessions. PVs were significantly decorrelated after learning when compared to the control data (Fig. 2c; Kruskal Wallis test, *H*(3) = 147.9, *p* < 0.001), yet no significant difference was detected in the mean rate change analysis (Fig. 2d; Kruskal Wallis test, *H*(3) = 6.6, *p* = 0.086), and only a difference between learning sessions and the shockCtrl, but not noShockCtrl, was observed in the place field correation analysis (Fig. 2e; Kruskal Wallis test, *H*(3) = 16.6, *p* < 0.001). Thus, despite a significant remapping at the population level, many cells did not remap, indicative of partial remapping.

Next, we asked whether remapping occurred across all arms of the maze. On the path leading to the choice point (shown in green in Fig. 2a), the lowest PV correlations were actually observed during the noShockCtrl sessions, indicating that rather than causing remapping, task learning actually stabilized the map on this part of the track (Fig S2a; Kruskal Wallis test, *H*(3) = 36.6, *p* < 0.001). In contrast, on the choice arms, robust remapping was observed during learning sessions relative to both control sessions (Fig. S2b; Kruskal Wallis test, *H*(3) = 165.2, *p* < 0.001). Examining the individual choice arms, we found that learning-induced remapping occurred primarily on the arms whose contingencies changed (i.e. those arms that were initially “safe” that switched to “shock”, or vice versa), as opposed to those that didn’t (i.e. the safeToSafe arm from Learning1 and the shockToShock arm from Learning2; Fig. S2c; Kruskal Wallis test, *H*(7) = 206.1, *p* < 0.001). Further, this analysis showed that during shockCtrl sessions, the spatial representations of the stable shocked arms (shockToShock) arms were more stable than those of the stable no shock arms (safeToSafe). Thus, task learning leads first to remapping, followed by stabilization occurring primarily at the aversive locations.

### Remapping occurs in a specific place cell population

Next, we wanted to identify remapping and non-remapping cells so that we could compare their properties. We combined the rate maps (excluding the “before choice” region of the track where no remapping was observed) of all place cells recorded in all 4 types of sessions, creating 2 sets of pre- and post-learning rate maps from which we calculated an average PV correlation value. We then performed a subtraction analysis, omitting each cell individually and recalculating the average pre- and post-learning PV correlation; the difference between this value and the original value was then taken as a measure of the cell’s contribution to remapping, with positive values indicating a contribution to remapping and negative values indicating a contribution to stability. Given that place cell firing rates can be modulated by novelty^32, 33^, we decided to use each cell’s maximal firing rate in combination with its remapping contribution to aid in identification of the remapping cell population. Plotting maximal firing rates against remapping contribution, it was evident that the majority of cells had relatively low firing rates and made very little contribution to either remapping (positive remapping contribution values) or stability (negative remapping contribution values). However, there were a large number of cells with relatively high firing rates and either large positive or large negative remapping contribution values. We ran *k*-means clustering on data (after first *z*-normalizing it) using 3 seed clusters, repeated 1000 times. Of the 735 place cells included in the analysis, only 2 cells did not consistently (< 95% of repetitions) sort into the same cluster (Fig. 3a). Further, the centroid locations did not move across repetitions, in contrast to those calculated using shuffled data (Fig. 3b), confirming the robustness of the clustering. Place cell populations recorded during the learning sessions had a significantly higher proportion classified as remapping cells compared to those recorded during control sessions (Fig. 3c; *p* = 0. 020), consistent with the greater remapping measured at the population level (see Fig. 2). In total, we identified 137 remapping, 182 stable cells, and 414 low firing rate cells (Fig. 3d-h, Fig. S3). This suggests that during episodic learning, distinct populations of place cells are responsible either for encoding the experience through remapping or maintaining the map of the physical environment.

**Figure 3.**
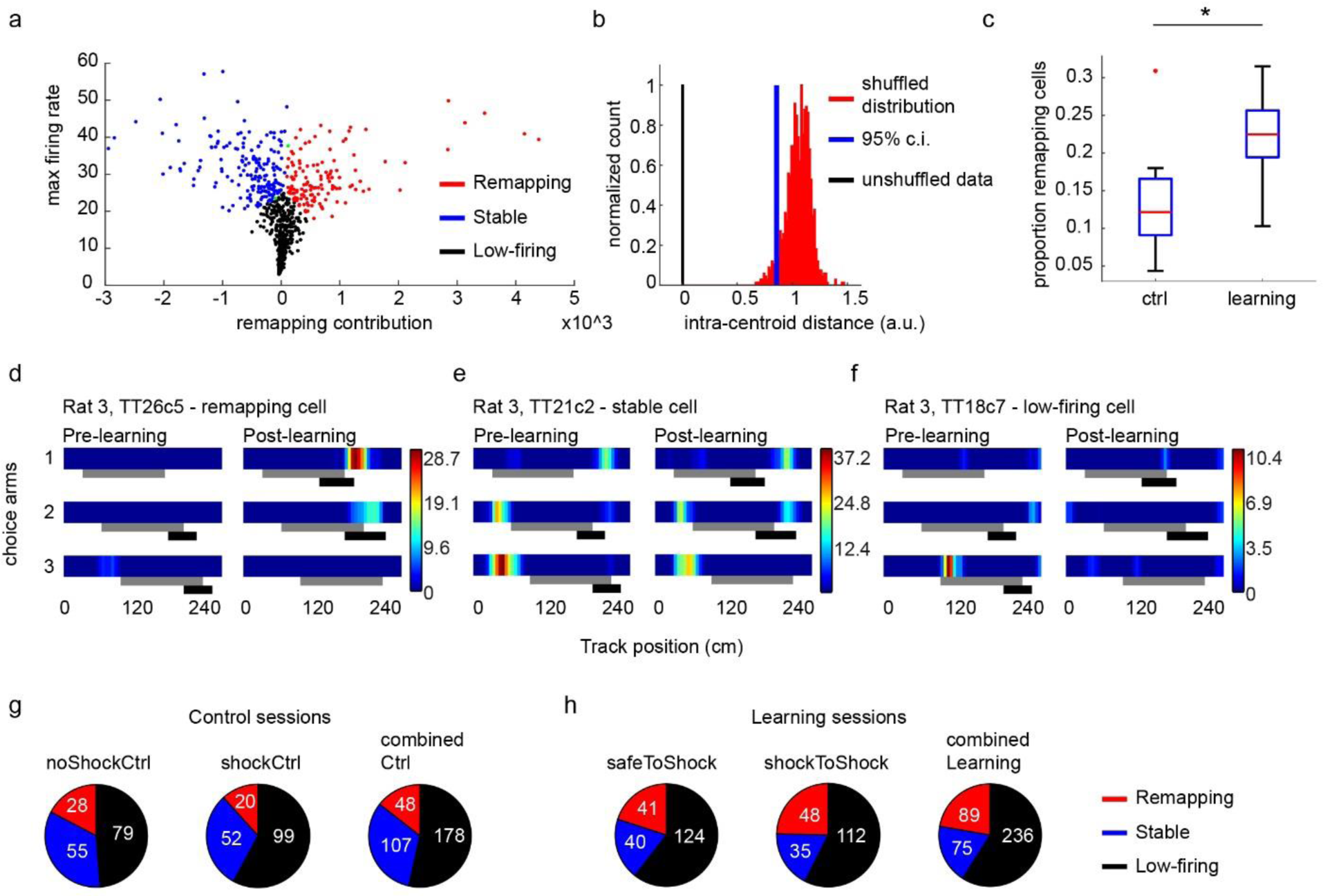
Identification of remapping, stable, and low-firing cell populations. (**a**) Each unit’s maximal firing rate plotted against its remapping contribution. *K*-means clustering of the z-normalized data using 3 seed clusters reliably partitioned the data (raw data is shown for display only). Remapping cells in red, stable cells in blue, low-firing cells in black, and the 2 cells which didn’t reliably sort into a cluster (i.e. less than 95% trials in a single cluster) in green. (**b**) The mean distance between cluster centroids on consecutive trials of *k*-means clustering was 0, indicating constistent clustering of the data. Randomly pairing firing rate and remapping contribution data prior to clustering created the distribution in red when repeated 1000 times. The left side of the 95% confidence interval is shown in blue. (**c**) The proportion of total cells per session which were classified as remapping cells. Wilcoxon rank sum test, *p* = 0.02. (**d**) An example of the choice arm rate maps for 1 remapping cell. The grey box under the linearized rate maps indicates the central portion of the choice arm, and the black box indicates the length of track over which the animal received the aversive eye-lid shock. (**e**) An example stable cell, and (**f**) an example low-firing cell, both recorded during the same shockToSafe session as the remapping cell recorded in (**d**). (**g**) Pie charts showing the number of each cell type identified during each type of ctrl session (left and middle) as well as combined across the two session types. (**h**) As in (**g**), but for the learning sessions.

### Remapping cell characteristics

We examined firing rates across the 3 cell types. Remapping cells initially had lower firing rates than stable cells, but their rates increased during the behavioral sessions, both during control (Figure 4a, c-e; Kruskal Wallis test, *H(*5) = 396.2, *p* < 0.001) and learning sessions (Figure 4b, c-e; Kruskal Wallis test, *H*(5) = 384.2, *p* < 0.001). Place fields expanded slightly in remapping cells in both the control and learning sessions, while those of stable cells did not change their scale (low-firing place fields contracted slightly only during the control sessions; Fig. S4; Wilcoxon signed-rank test, ctrl sessions: remapping, *Z* =2.09, *p* = 0.037; stable, *Z* = −0.319, *p* = 0.754; low-firing, *Z* = −2.84, *p* = 0.0045; learning sessions: remapping, *Z* = 2.07, *p* = 0.039; stable, *Z* = −1.91, *p* = 0.057; low-firing, *Z* = 0.950, *p* = 0.3419).

**Figure 4.**
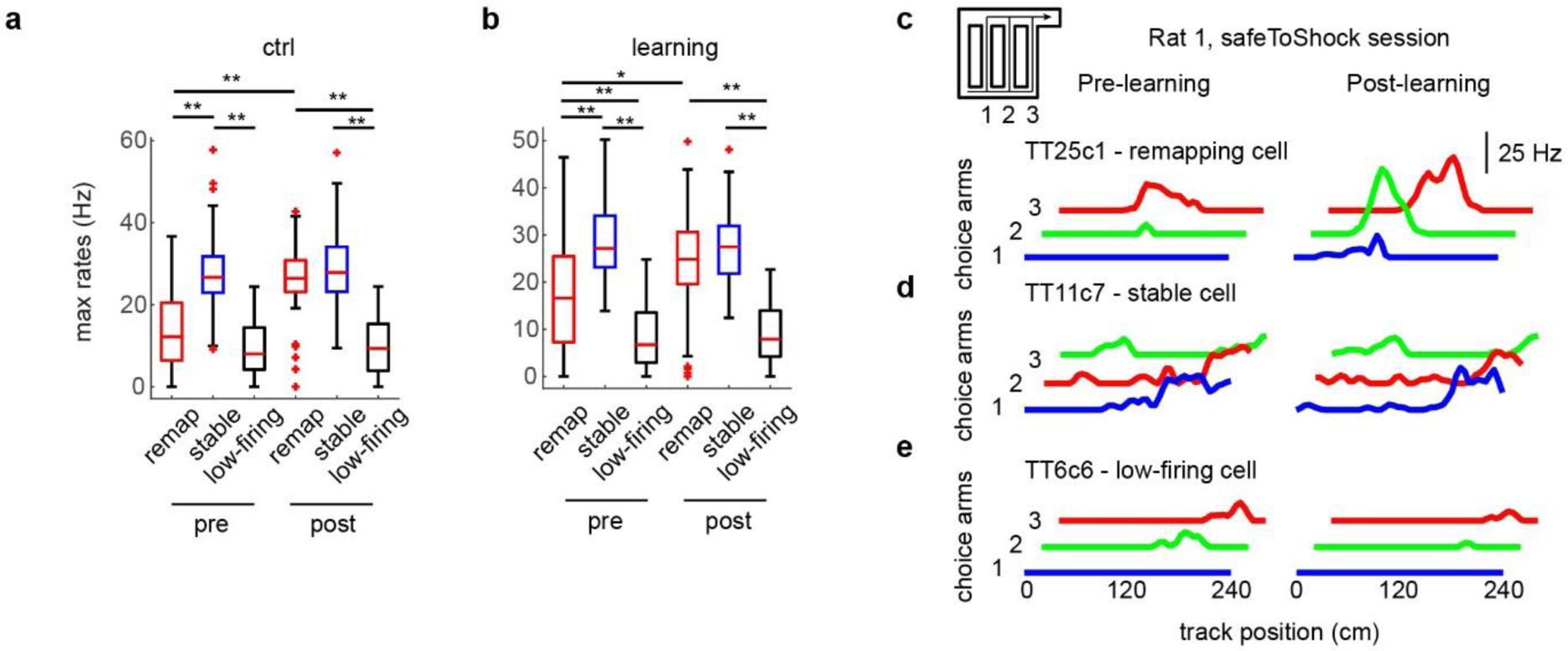
Maximum firing rates of remapping and stable cells. (**a**) Maximum firing rates of all cell types during ctrl sessions recorded during the beginning (baseline) and end (post) of the sessions (*n* = 48 (remapping cells), 107 (stable cells), 178 (low firing cells); Kruskal Wallis test, *H*(5) = 396.2, *p* < 0.001). (**b**) Maximum firing rates of all cell types during during learning sessions recorded during the beginning (baseline) and end (post) of the sessions (*n* = 89 (remapping cells), 75 (stable cells), 236 (low firing cells); Kruskal Wallis test, *H*(5) = 384.2, *p* < 0.001). * and ** denote significance levels p < 0.05 and p < 0.001 respectively. (**c**) Example choice arm firing rate plots from a remapping cell recorded during a safeToShock session. Inset, layout of maze with locations of choice arms 1-3. (**d**) Example rate maps from a stable cell and (**e**) a low-firing cell recorded from the same animal as (**c**).

We wondered whether remapping might be induced by shock-mediated excitation of remapping cells. To investigate this, we constructed peri-event time histograms (PETHs) centered on the shock times; it should be noted that these PETHs could only incorporate spikes from cells that happened to have place fields overlapping the shock locations, and therefore, many place cells did not contribute spikes to this analysis. Because we varied shock-intensity, reducing it as animals learned the new contingencies (see Methods), we selected the 10 highest-intensity shocks from each session for this analysis. Both remapping and non-remapping populations displayed some shock-induced suppression relative to low-firing cells (Fig. S5a-e; *p* < 0.5 for both populations), ruling out shock-induced modulation as an underlying mechanism for remapping.

### Remapping cell preference for earlier theta phases, and enhancement during encoding

Encoding of spatial information is thought to occur on the ascending wave of theta, when entorhinal inputs, providing real-time sensory information, dominate. Recall, on the other hand, is thought to occur on the descending wave when CA3 inputs, providing information stored within its recurrent connections, are strongest^21, 23, 34^. We therefore hypothesized that remapping cell spiking might display a theta phase preference shifted towards the ascending wave of theta. In control sessions, the theta phase preference of remapping cells was slightly shifted in the opposite direction, towards the descending wave, relative to low-firing cells (Fig. 5a, c, d; bootstrap statistics shown in Fig. S6a). In the learning sessions, this shift was substantially larger (Fig. 5b-d; Fig. S6d, g). Both the low-firing and stable cells shifted their firing to the descending phase as well, but not to the same degree as remapping cells (Fig. S6d-i). Comparing the ratio of spikes fired on the descending wave relative to the ascending wave between the 3 cell types confirmed these finding (Fig. 5d; Fig. S6a-i).

**Figure 5.**
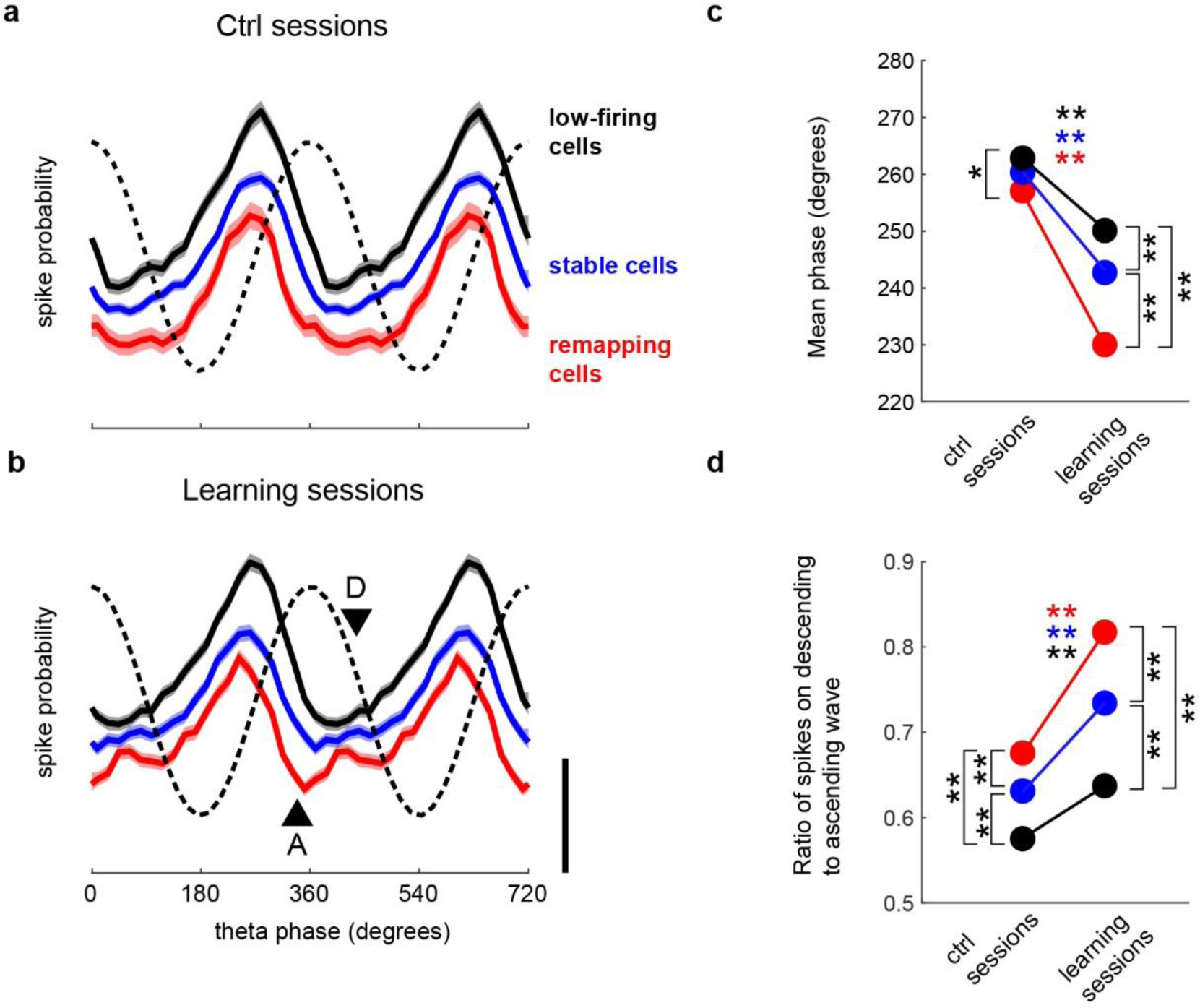
Shifted theta phase preference of remapping cells. (**a**, **b**) Theta phase distributions of remapping, stable, and low-firing populations in control (**a**; *n* = 45345 (remapping), 160427 (stable), and 74705 (low-firing) spikes) and learning sessions (**b**; *n* = 105162 (remapping), 111110 (stable), and 106109 (low-firing) spikes). Black triangles indicate decreased preference of remapped field spikes for the ascending wave (A)/peak and increased preference for the descending wave of theta (D); these are indicated for illustrative purposes only, statistical analysis of ratio of spikes fired at D vs. A shown in (**d**). Vertical scale bar indicates a difference in proportion of 0.02. (**c**) The mean theta phases of remapping, stable, and low-firing cell spikes in control and learning sessions. Bootstrap statistics shown in Fig. S6. (**d**) Mean ratio of spikes fired on the descending wave to the ascending wave for all 3 cell populations. Bootstrap statistics shown in Fig. S6. Colored stars indicate a within cell-type difference across control and learning sessions. Black stars indicate differences between cell types. of remapped field (blue) or stable field (red) mean phase relative to baseline values (i.e. within group). Black stars aligned with vertical lines indicate significant difference from non-remapping cells during the same behavioral epoch. Statistics calculated as in (**c**). * and ** denote significance levels p < 0.05 and p < 0.001, respectively.

Because CA3 inputs are thought to be strongest on the descending wave of theta ^23^, we wondered if the shift of remapping cell spikes towards the descending wave of theta was being driven by CA3 inputs. CA3 inputs have previously been shown to drive increases in slow (30-50 Hz) gamma power^21, 23^, and we confirmed that slow gamma power was theta-modulated and greatest on the descending wave of theta (Fig. 6a, b). We wondered whether the shift in remapping cell theta phase preference during learning sessions was being driven specifically during episodes of high slow gamma power, which would implicate CA3. Indeed, we found that remapping cell spikes occurred at earlier theta phases during periods of elevated slow gamma power compared to periods of reduced power (Fig. 6d; Fig. S7a); while the shift in mean theta phase did not reach significance (Fig. 6e; Fig. S7b), the overall flattening of the phase distribution was also evident as a decrease in the mean resultant length of the distribution (Fig. 6f; Fig. S7c). Further, the shift during elevated slow gamma power was greater after the contingency change, suggesting it was driven by learning (Fig. 6g-j; Fig. S7d-f). Thus, the balance of inputs to remapping cells shifts towards CA3 during learning.

**Figure 6.**
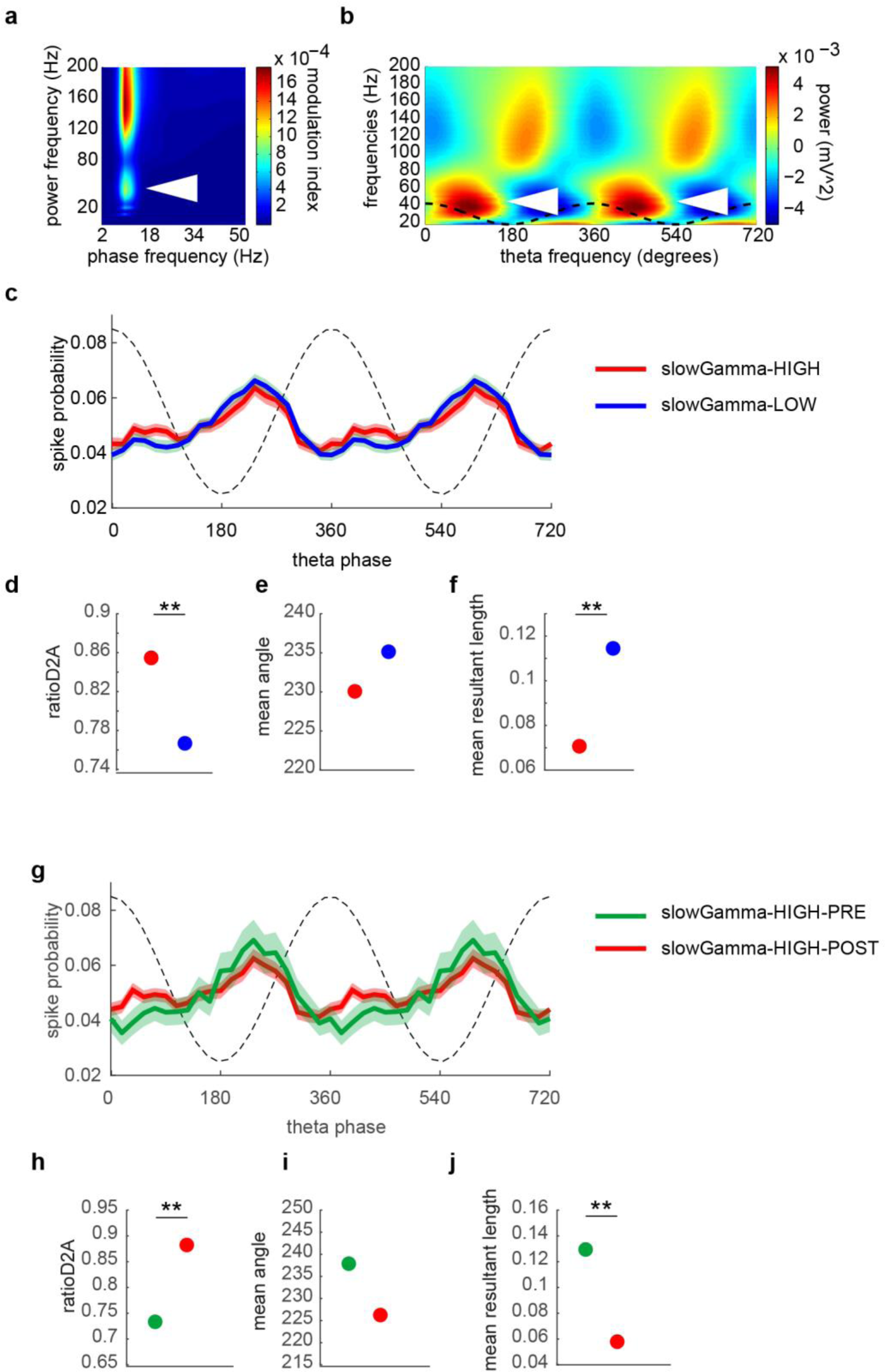
Shifted theta phase preference in remapping cells occurs during periods of elevated slow gamma power. (**a**) LFP phase-amplitude comodulogram, averaged across all tetrodes with place cells. (**b**) Gamma amplitude-theta phase modulation plot, same tetrodes as (**a**). Theta phase-modulated slow gamma power in (**a**) and (**b**) is indicated by white arrows. (**c**) Theta phase distribution of remapping cell spikes during periods of elevated (red, HIGH) or reduced (blue, LOW) slow gamma power (*n* = 30132 spikes (HIGH), 33031 spikes (LOW)). (**d**) The ratio of remapping cell spikes fired on the descending versus ascending wave of theta during elevated (red) or reduced (blue) slow gamma power. Bootstrap statistics shown in Fig. S7. (**e**) The mean phase of remapping cell spikes during elevated (red) or reduced (blue) slow gamma power. Bootstrap statistics shown in Fig. S7. (**f**) The mean resultant length of the remapping cell spike-phase distribution during elevated (red) or reduced (blue) slow gamma power. Bootstrap statistics shown in Fig. S7. (**g**) Theta phase distribution of remapping cell spikes during periods of elevated slow gamma power during learning sessions before (red) and after (green) the contingency change (*n* = 5399 spikes (before), 24733 (after)). (**h**) The ratio of remapping cell spikes fired on the descending versus ascending wave of theta during learning sessions before (red) and after (green) the contingency change. Bootstrap statistics shown in Fig. S7. (**i**) The mean phase of remapping cell spikes during learning sessions before (red) and after (green) the contingency change. Bootstrap statistics shown in Fig. S7. (**j**) The mean resultant length of the remapping cell spike-phase distribution during learning sessions before (red) and after (green) the contingency change. Bootstrap statistics shown in Fig. S7. ** denotes significance level of p < 0.001.

### Remapping cells increase their replay participation during learning

Awake replay of hippocampal place cell sequences has been proposed to function as a consolidation mechanism to stabilize plasticity and link place cell networks following learning^15, 35^. A recent study showed that in a novel spatial environment, new spatial information may be integrated in the hippocampus through replay of place-cell sequences during sharp wave ripples (SPW-Rs) by a subset of “plastic” place cells^24^, suggesting the hypothesis that contextual information might be incorporated through enhanced replay participation by remapping cells. We detected transient elevations of the multi-unit firing rate co-occurring with SPW-Rs during breaks in the animal’s running. Across learning, both remapping and low-firing, but not stable, cells increased their ripple-centered firing rates (Fig. 7a, b; Wilcoxon signed-rank test, remapping cells: *p* < 0.001; stable cells: *p* = 0.069; low-firing cells, *p* < 0.001); however, the increase in the remapping cell group was significantly greater than the low-firing group (Kruskal Wallis test, *H*(2) = 15.4, *p* < 0.001; Fig. 7b).

**Fig. 7.**
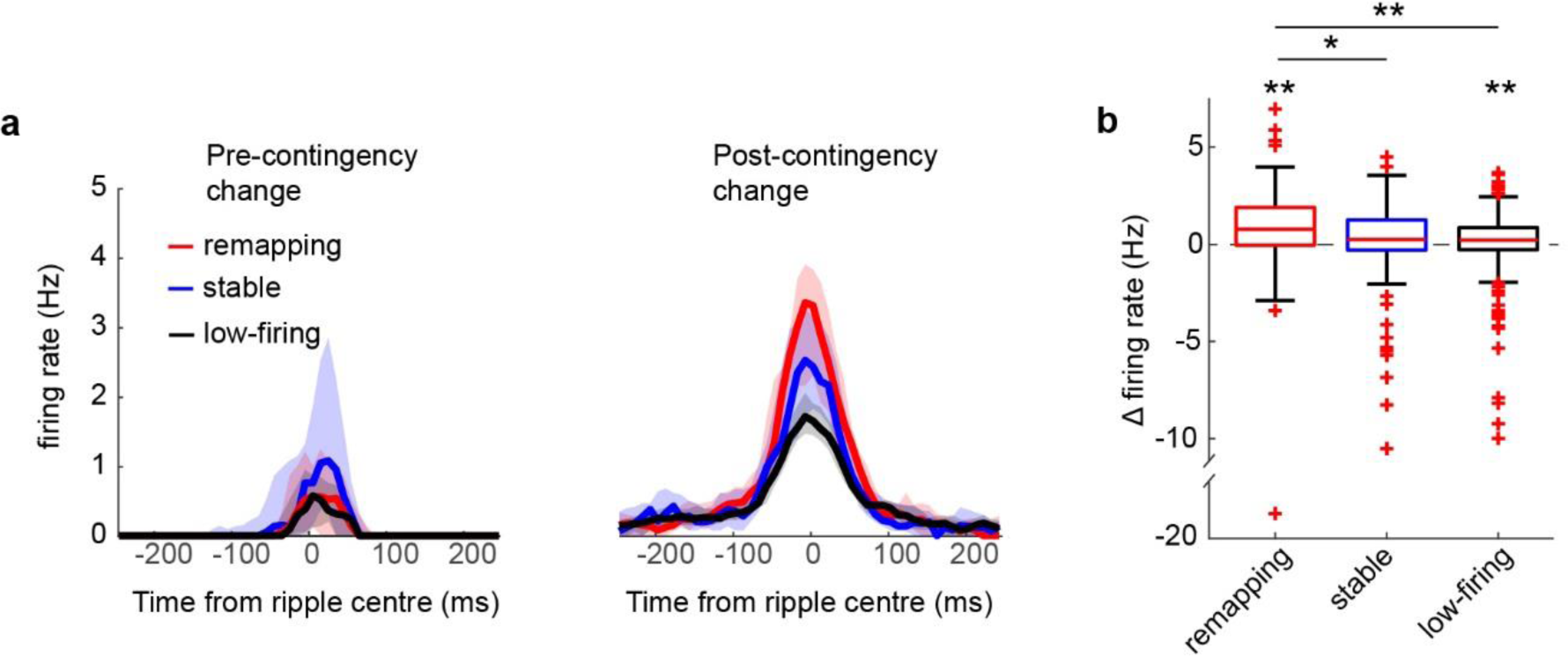
Ripple-centered activity of remapping place cells. (**a**) Ripple-centered firing rate during learning sessions of remapping (red; *n* = 89), stable (blue; *n* = 75) and low-firing (blue; *n* = 236) cells pre- (left) and post- (right) contingency change. Shaded area represents 95% confidence interval. (**b**) Change in firing rate between post-learning and pre-contingency change epochs (Wilcoxon signed-rank test, remapping cells: *p* < 0.001; stable cells: *p* = 0.069; low-firing cells, *p* < 0.001) for each cell type, as well as difference in change between cell types (Kruskal Wallis test, *H*(2) = 15.4, *p* < 0.001). * and ** denote significance levels of *p* < 0.05 and *p* < 0.001, respectively.

To investigate replay directly, we employed Bayesian decoding^36^ (Fig. 8a) of spikes fired during transient elevations in the population firing rate. The majority of these events occurred at the reward location (Fig. S8a), with a slight bias towards reverse replay (Binomial test: *p* = 0.033; Fig. S8b), though no bias towards encoding of the next choice during forward replay (*p* = 0.492; Fig. S8c) nor the previous choice during reverse replay (*p* = 0.876; Fig. S8d), nor any bias towards encoding either shock or safe arms (i.e. no significant deviation from a 2:1 ratio of shock to safe events, *p* = 1; Fig. S8e).

**Fig. 8.**
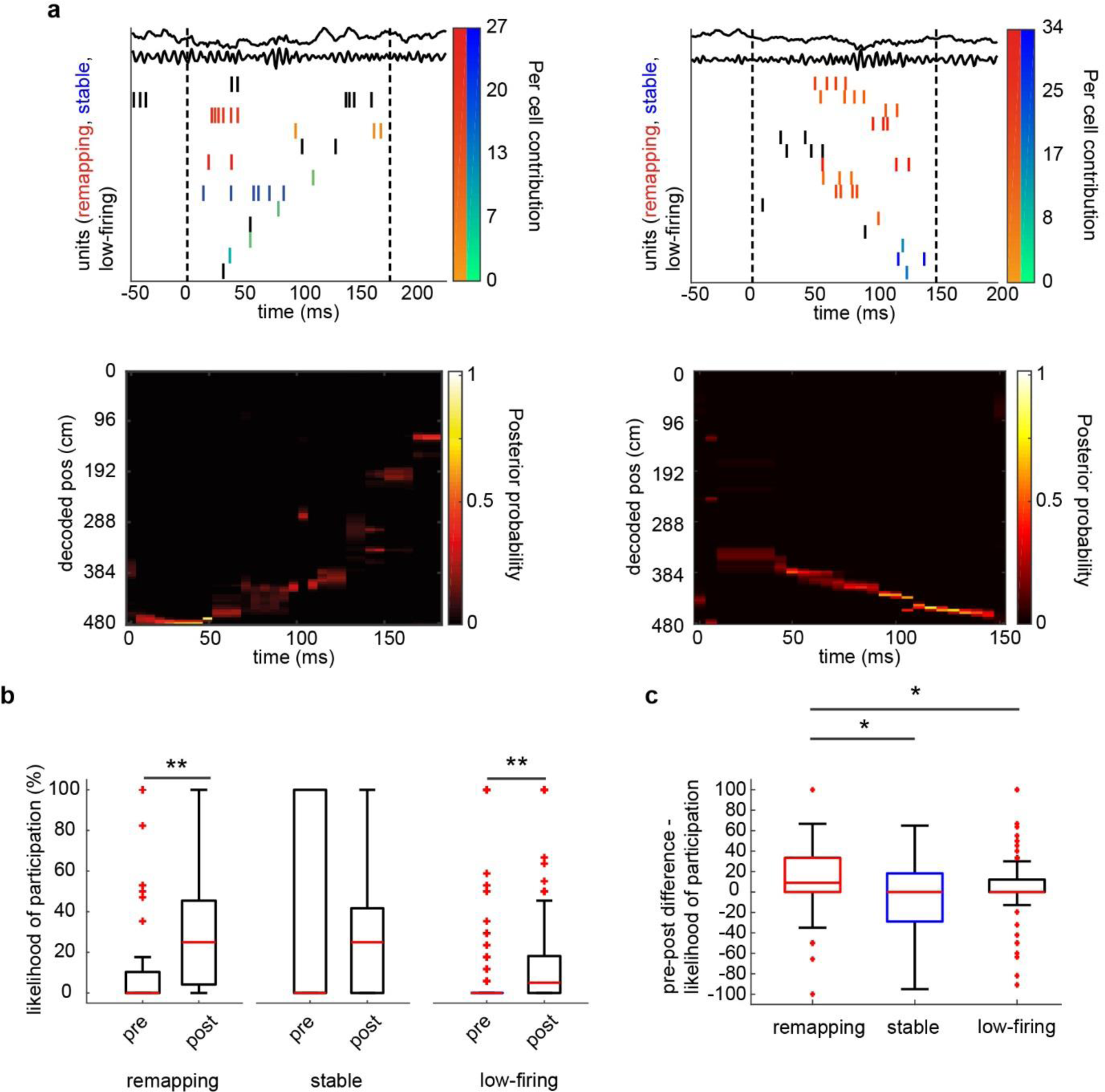
Awake replay of remapping place cells. (**a**) Example replay events from pre-contingency change (left) and post-learning (right) epochs showing the raw and ripple band filtered LFP and spikes color-coded by cell identity (remapping – autumn colors; non-remapping – winter colors) and each cell’s Per Cell Contribution (PCC) score, and ordered according to their location of peak firing (top; vertical dashed lines show the temporal borders of the event) and heat maps showing the posterior probabilities calculated using Bayesian decoding (bottom). (**b**) Proportion of replay events in which cells were active during pre-contingency change and post-learning epochs (Wilcoxon signed-rank test, remapping: *Z* = −4.79, *p* < 0.001; stable: *Z* = 1.52, *p* = 0.128; low-firing: *Z* = −6.470, *p* < 0.001; Fig. 8d). (**c**) Pre to post change in proportion of events in which cells participated (i.e. fired at least 1 spike; Kruskal Wallis test, *H*(2) =, *p* = 0.0147). * and ** denote significance levels of *p* < 0.05 and *p* < 0.001, respectively.

To assess the degree to which the different cell populations changed their participation in replay events across learning, we calculated the proportion of replay events in which each cell was active (i.e. fired at least 1 spike). Both remapping and low-firing cells, but not stable cells, increased the proportion of replay events in which they were active across learning (Fig. 8b; Wilcoxon signed-rank test, remapping: *Z* = −4.79, *p* < 0.001; stable: *Z* = 1.52, *p* = 0.128; low-firing: *Z* = −6.470, *p* < 0.001); the increase was most pronounced in remapping cells (Fig. 8c; Kruskal Wallis test, *H*(2) = 8.44, *p* = 0.0147). However, when examining only events within which a cell participated, we found no differences across learning in either the number of spikes fired per event (Fig. S9a, b; within groups, Wilcoxon signed-rank test, remapping: *Z* = 0.724, *p* = 0.469; stable: *Z* = 0.577, *p* = 0.577; low-firing: *Z* = 0.252, *p* = 0.801; across groups, Kruskal Wallis test, *H*(2) = 0.323, *p* = 0.851), nor in Per Cell Contribution score (a measure of a cell’s contribution to the correlation between time and decoded position within the replay event)^24^ (Fig. S9c, d). These data suggest that integration of new information into the spatial map involves an increase in the number of events in which remapping cells are active, rather than a change in their spiking behavior within those events.

## DISCUSSION

Our data show that in a familiar environment, changes in behavioral and emotional context initiate a number of changes indicative of episodic memory encoding in a subset of CA1 place cells. We find that partial remapping^17, 18^ is not simply a stochastic process in which some proportion of a homogeneous place cell population remaps, but rather results from remapping in a predefined cell population characterized by their low pre-learning firing rates and replay participation scores, and shifted spike-timing relative to the theta oscillation. With learning, we observe a further shift in remapping cells’ spike timing to even earlier phases of theta as well as increases in their firing rates and participation in awake replay events. The modulation of this shifted spike timing by slow-gamma power supports a role for CA3 inputs in driving remapping^21, 23^, as does the place field size increase with learning^37^. Together, our data identify memory encoding mechanisms which allow for reorganization in a place cell sub-population while also allowing stable spatial coding to be maintained in a separate population.

One interesting aspect of our study is that we identified remapping cells not just during learning, but also during control sessions when our behavioral data indicated no additional learning took place. Importantly, there were significantly more remapping cells identified during learning, indicating that remapping can be used as a marker for learning. But the question remains, why is there any remapping during control sessions? If our interpretation that remapping cells are pre-selected is correct, then perhaps it is less surprising that in a population of cells predisposed to remap, some remapping will occur even in the absence of obvious contextual changes. In other words, a population with features that endow it with the ability to remap may have a harder time remaining stable than a population that is particularly suited for stability. Another possibility is that some of these remapping cells respond to changes in the environment that are neither relevant specifically to the task nor apparent to the experimenter, such as subtle changes in odor over the course of a recording.

In addition to the remapping and stable cell populations, we also identified a third cell population, which we termed low-firing cells. Owing to their low-firing rates, these cells made minimal contributions to the stability of the population-wide spatial representation. It could be argued that these cells are simply low-firing members of the remapping and/or stable populations; however, both remapping and stable populations had theta phase preferences that were shifted to earlier phases relative to the low-firing population, even before learning, suggesting they do form a distinct population.

What is not clear is whether these populations remain segregated at all times, or whether individual cells can leave one population and join another. It would seem that some fluidity is required, otherwise, once the information encoded by remapping cells is consolidated, how can remapping repeat the next time new information must be learned? Imaging studies have shown quite clearly that there is substantial turnover of the active population in CA1 over days ^38^, suggesting a possible cycle whereby low-firing cells become remapping cells first before later stabilizing, and then ultimately turning-over, becoming low-firing rate cells again. Future studies tracking these cells across multiple days will be required to clarify this issue.

### Inputs driving remapping

Encoding and retrieval of memory are thought to occur on different phases of theta as a way of preventing interference between the two processes. In this view, encoding occurs on the ascending wave and peak of theta when entorhinal inputs and the real-time, real-world information they provide, dominate, while retrieval occurs on the descending wave when inputs from CA3, transmitting mnemonic information stored within the recurrent network there, are strongest^19^. In support of this model, inhibition of CA1 place cell activity immediately after the theta peak or trough causes modest enhancements in encoding or retrieval, respectively, in a spatial working memory task^39^. Further, in a novel environment, place cells have been shown to shift their firing away from the trough of theta towards the ascending wave^40, 41^. We therefore predicted that remapping cells would have theta phase preferences shifted towards the peak of theta and away from the descending wave. However, we observed the opposite – relative to low-firing cells before learning, and relative to both low-firing and stable cells post-learning, remapping cells fired more on the descending phase, when CA3 inputs are strongest^21, 23, 34^, and less on the ascending phase, when entorhinal inputs dominate^23, 34^. Interestingly, all 3 types of place cells displayed a shift towards the descending wave with learning, though this shift was greatest in remapping cells. Recently, a place cell study of engram cells ^42^ in the mouse hippocampus showed that when engram cells fire bursts of action potentials, they tend to do so on the descending wave of theta ^43^, lending support to our findings and suggesting that the remapping cell population may overlap with that of engram cells.

In remapping cells after learning, the shift was most pronounced during episodes of elevated slow-gamma power when CA3 inputs are dominant ^21, 23, 34^, suggesting it resulted from a shift in the balance of inputs onto remapping cells towards CA3 and away from entorhinal cortex. This interpretation is further supported by a report that lesion of the direct medial entorhinal layer III inputs to CA1 leads to increased spatial scale^37^, which we observed in remapping cells. Furthermore, an important role for CA3 in our behavioral task is supported by previous studies which found that CA3-CA1 LTP is necessary for contextual fear learning^44^ and leads to partial remapping in CA1^45^. Together, this suggests that distinct from the learning of spatial features, episodic memory encoding engages a CA3-CA1 circuit which modulates activity more specifically in remapping cells.

Previous work has shown that optogenetic inhibition of CA1 place cells can produce remapping, suggesting a functional role of inhibition in this process^46^. Further, suppression of CA1 excitability by aversive sensory stimulation has previously been reported^47–49^, and been shown to be due mainly to septal cholinergic inputs^50^. These inputs activate somatostatin-positive interneurons, and activity in these cells is necessary for contextual fear learning^51^. Here, we observe shock-induced suppression in both remapping and stable populations, suggesting this suppression is not specific to cells involved in encoding new information. Further, many remapping cells had fields not overlapping with the shock zone, and thus would not have been suppressed by the shock. Nevertheless, the possibility remains that this suppression may have been required specifically in those cells with fields overlapping the shock-zone, as this suppression is thought to prevent sensory features at the time of memory induction from interfering with the encoding of context by the hippocampus^51, 52^. Further, acetylcholine release has also been reported to have more long-lasting excitatory effects in CA1 after the initial suppression^50^, suggesting that acetylcholine might be acting not only on cells with fields at the shock zones, but also on cells with more distal fields. Given recent studies revealing molecular heterogeneity in hippocampal neurons^53^, another possibility is that the different cell populations may exhibit unique expression patterns, such as for acetylcholine receptors, or distinct patterns of connectivity.

### Replay as a memory consolidation mechanism occurring in remapping cells

Previous studies have shown that in reward learning, cells whose fields become clustered around the reward zones increase their firing during SWRs^15^. However, because reward increases firing rates during SWRs, and SWRs tend to involve cells that have place fields at the reward zone^25^, it was not clear whether the enhanced contributions of reward zone cells was due to a unique role for these cells in replay, or simply a by-product of their place field locations. Our task, in which remapping and non-remapping cell place fields were distributed along the track arms, circumvented this limitation, and our data show that cells involved in memory encoding increase their contributions during learning, whereas stable, high-firing rate cells do not.

While there is ample evidence that replay episodes are involved in planning behavior^3^, we didn’t observe any bias of replay towards the immediate future path. However, a recent report showed that after rats were shocked at one end of a linear track, they subsequently replayed the “shock-zone” even though they never re-entered it^18^, suggesting that replay episodes in our task might be involved in both planning where and where not to go, making observation of planning during replay difficult. The lack of any significant bias towards either future or previous paths is also consistent with observations that replay may reflect all physically available trajectories, which was suggested to provide a method for learning and maintaining the cognitive map^54^.

### Conclusion

Our identification and examination of remapping cells shows that increased coordination by CA3 and enhanced replay in these cells may facilitate the generation and consolidation of plasticity in the memory encoding place cell network, thereby linking a spatially restricted experience to a representation of the larger environment in which it occurred. The enhancement of replay contribution in remapping cells suggests the existence of a mechanism to select those cells with the emergent coding properties that best support adaptive behavior for participation in replay. That these processes occur in a specific cell population, while a separate population of place cells exhibit stable spatial coding, demonstrates a mechanism whereby the hippocampus can both maintain a stable spatial representation of the environment while incorporating changing features of experiences that occur within that environment.

### Author Contributions

J.O. and J.P.J. designed the experiments and wrote the paper. J.O. and S.A.S. collected data. J.O. analyzed data.

## Acknowledgments

We thank Mark Turnbull for providing experimental support and Thomas McHugh, Shigeyoshi Fujisawa and Charles Yokoyama for providing comments on earlier versions of the manuscript.

## Experimental Procedures

A total of 4 Long Evans rats were used. Animals were 6-11 months of age at the time of data collection. Experimental procedures were approved by the Animal Care and Use Committees of the RIKEN Brain Science Institute. Animals were food deprived to ∼85-90% of their baseline weight, and trained to nose poke for food reward (sweetened condensed milk diluted with an equal amount of water) on the track. Typically, animals required 3 to 4 sessions to reach > 100 nose pokes. We then began training the animals to run laps on the full track for food reward. Rats ran an equal proportion of forced and free choice trials during training. Once animals could run > 100 trials, we implanted with electrode arrays targeting dorsal hippocampus, as well as basolateral amygdala (BLA) and lateral entorhinal cortex (LEC) (BLA and LEC data is not presented in this manuscript).

### Surgery

Rats were anesthetized with .5-1.5 % isoflurane. Two craniotomies were made over dHPC (4.2 mm posterior from Bregma, ±3 mm lateral from midline). Periorbital shock wires (stainless steel, insulated 0.003in/0.055in diameter, A-M Systems cat. # 791000) were implanted beneath the skin of each eyelid. The electrode array, containing 26 to 28 tetrodes targeted to each hippocampus (13 to 14 tetrodes per hemisphere), was implanted on to the surface of the cortex and electrodes turned 750 µm into the brain. Two bone screws attached to the skull served as ground and reference. Tetrodes (nichrome, ¼ Hard Pac coating, 0.0005 in diameter, Kanthal item # PF000591) were gold-plated to < 150 kΩ prior to implantation. Tetrodes were lowered to dorsal CA1 over two weeks, and rats continued to run daily training sessions on the track. Once tetrodes were in the CA1 cell body layer, data collection commenced. Data was acquired using a Neuralynx Digital Lynx acquisition system. Amimals were tracked using custom LEDs imaged by an overhead camera at a frame rate of 30 fps.

### Behavior

A sequence of four sessions was run in each animal. In all sessions, the animal left the start/reward zone and ran to a small reward zone where it nosed poke for a small food reward (20 µl sweetened condensed milk). This triggered a door to close behind it, and another door to open in front of it. It then proceeded to run to a choice point where it could choose one of three arms in free choice trials, or a single arm due to closing of the other two arms during forced choice trials, on which to run back to the main reward area where it received a large food reward (100 µl sweetened condensed milk). It was then held in the start/reward zone for 10 seconds (by closing of the door behind it) before commencement of the next trial.

All 4 session types were composed of blocks of free choice trials, forced choice trials (which always alternated between the 3 choice arms – i.e. 1-2-3-1-2-3 etc.), and combined sequences of forced and free choice trials. These combined sequences were used to probe the animal’s memory/preference for the “safe” arm while forcing it to occasionally sample the shock arms; these sequences took the form of 2 free choice trials followed by a forced choice trial on one of the shock arms, then 2 more free choice trials followed by a forced choice trial on the other shock arm. To construct maps for remapping analysis (and other analyses restricted to pre- and post-learning epochs) we recorded pre- and post-learning baselines consisting of 4 trials on each arm; these could be recorded using 12 forced choice trials (noShockCtrl: both baselines; safeToShock: pre-learning baselines) or 4 repeats of the free choice – forced choice sequence (using the final 4 free choice trials on the “safe” arm and all 8 (4 × 2) forced choice trials on the shock arms; safeToShock: pre-learning baselines; shockCtrl and shockToShock: all baselines). In some cases, we recorded additional trials after the post-learning baseline to be used for the awake replay analyses. To probe memory for the previous sessions contingencies, each session began with 5-10 free choice trials. These probe trials also allowed us to determine a “least preferred” arm (i.e. the arm chosen least frequently by the animal), which would then be used as the subsequent “safe” arm in the next contingency change; using the “least preferred” arm allowed us to ensure that subsequent preference for this arm reflected new learning rather than expression of a previously held preference. If there was a tie between 2 arms for the designation of “least preferred” (i.e. the animal chose 1 arm 5 times and the other arms 0 times), we broke the tie by examining the previous session’s behavioral data. A change in contingency was always immediately followed by forced choice trials to ensure that the animal sampled all arms (pilot experiments indicated that animals were resistant to change their choice behavior away from a previous safe arm to a new safe arm if they weren’t first forced to sample the new safe arm). Shocks were delivered as bilateral 1 second trains of 1 ms pulses delivered at a frequency of 7Hz (i.e. 7 pulses total). Because of differences in shock sensitivity and behavior between animals, the experimenter had to occasionally adjust shock intensity as well as introduce additional forced choice trials in order to ensure that the animal learned the new contingencies. Once the contingencies were learned (20 trials above 75% correct choices) and the post-learning baseline was to be recorded, shock intensity had to be reduced to a level where the animal would willingly run through the shock on forced choice trials but would maintain their preference for the shock arm on free choice trials. Shock intensities ranged from 0.4 – 1.5 mA. The total number of trials run was also determined by how quickly the animal learned the contingencies in the learning sessions, and in all sessions the experimenter had to judge the animal’s level of motivation when deciding to run more trials or end the session. There were no significant differences in the number of trials run between the different session types (One-way ANOVA, *p* = 0.89).

### Spike sorting

Spike sorting was performed manually with MClust (A. David Redish, University of Minnesota, Minneapolis, MN) using two-dimensional projections of waveform amplitudes and energies, and autocorrelation and crosscorrelation functions as additional separation tools and separation criteria. Excitatory cells were distinguished from interneurons by spike width and average rate. L-ratio (median = 0.05), isolation distance (median = 27.0; Schmitzer-Torbert et al., 2005) and percentage of inter-spike intervals < 2 ms (median == 0) were used as metrics to assess cluster quality.

### Histology

In rats #2 and #3, marking lesions were made using 20 µA of anodal current for 10 seconds; in animals #1 and #4, no marking lesions were made. Animals were transcardially perfused with phosphate-buffered saline followed by 4% paraformaldehyde (PFA), brains cryoprotected in 30% sucrose/4% PFA, and frozen slices of 20 µm (animals #1 and #4) or 40 µm (animals #2 and #3) were cut using a cryostat, and stained using either NeuroTrace 530/615 Red Fluorescent Nissl Stain (ThermoFisher) or DAPI (Sigma) (Fig.S1).

### Analysis

All data analyses were performed in Matlab.

### Task performance (Figure 1)

Performance in the task was calculated using a moving average (5 trials total including the current trial, the two previous and two next trials; Fig. S1b). In Figure 1g-i, the “pre” percent correct was calculated using the first 5 free choice trials of the session and the “post” percent correct was calculated using the last 20 free choice trials of the session.

### Place fields

Positional data was extracted from the video files, smoothed, and restricted to times when the animal was moving at faster than 5 cm/s for at least 1.5 seconds (durations of less than 0.3 sec below 5 cm/s within these runs were permitted). The 2-dimensional positional data was then linearized (as outlined in Figure 2a). Each of the 4 arms (1 preChoice arm, and 3 choiceArms) was divided into 50 spatial bins (4.8 cm/bin), and spikes from individual units were assigned to those bins using linear interpolation. Spike counts were then divided by occupancy at each spatial bin to produce firing rates, and the resulting rate maps were smoothed. A unit with a peak firing rate ≥ 3 Hz and spatial information ≥ 0.3 bits/spike was classified as a place cell.

### Remapping (Fig. 2, Supplementary Fig. 2)

Population vector correlations, place field correlations, and mean rate differences were calculated to quantify remapping. In the population vector analysis, pairs of vectors containing the firing rate of each place cell at a given bin before and after learning were constructed and the Pearson’s correlation calculated. Thus, for each session, there were 200 values of Pearson’s r (50 bins per arm multiplied by 4 arms) contributed for subsequent statistical analysis. In the place field correlation analysis, Pearson’s correlation was calculated for pairs of pre- and post-rate maps for individual place fields. In the mean rate difference analysis, the mean rate of the pre rate map was subtracted from the mean rate of the post rate map, and the absolute of this value was divided by the sum of the two mean rates (thus a value of 1 indicates zero firing in either the pre or post epoch, and a value of 0 indicates identical mean rates in the two epochs).

### Identifying remapping cells (Fig. 3)

Rate maps from each recorded cell, omitting the before choice portion of the path were combined to create pre- and post-rate maps of the entire population. For each cell, the contribution of each individual rate map to decorrelation of the PVs (i.e. remapping) was calculated as follows: first, the population of PV correlations between all pre- and post-learning rate maps were calculated, which was then used to calculate a mean PV correlation value; second, the PV correlations were recalculated N times, where N refers to the number of place cells, each time omitting the *n*th cell’s rate map, and these were used to generate N mean PV correlation values; third, the mean value generated using all rate maps was subtracted from the mean values generated from the partial sets of rate maps to generate a remapping contribution value for each rate map (e.g. a rate map demonstrating remapping remapping will generate a larger score because the mean PV correlation will be higher when that cell is omitted than when it is included). We then calculated the maximum in-field firing rate for each cell. Both the remapping contribution and the firing rate data were *z*-normalized. We then ran a *k*-means clustering algorithm (*kmeans*, MATLAB) on the data using 3 seed clusters; the clustering was repeated 1000 times, each time with the first seed being chosen at random from the data. Cells were included in 1 of the 3 clusters if it was assigned that cluster on greater than 95% of trials. The distances between each cluster’s centroids across consecutive trials of clustering was also calculated, and compared to a distribution calculated by repeating the clustering 1000 times on shuffled data. From this shuffled data, a 95% confidence interval was calculated; intra-centroid distance for the real data fell to the left of the confidence interval indicates that our clustering was consistent across trials, confirming the validity of our approach.

### Statistical significance of differences in shock-induced suppression of firing (Fig. S5)

To assess the shock-induced suppression of remapping and stable populations, we first normalized the population firing rates in the 1 sec after shocks by the mean firing rates and standard deviation in the 0.5 seconds preceding the shock. We then subtracted the mean normalized firing rates in that 1 second window from the low-firing population mean normalized firing rate. A control distribution was then constructed by creating 2 control populations with spikes randomly drawn from the remapping (or stable) and low-firing populations (but with each control population having the number of spikes as the real populations), then calculating the firing rate differences between these shuffled populations, and repeating the process 1000 times. A confidence interval was then created from this control distribution; the real differences for both remapping and stable cells fell to the left of the 95% confidence interval, indicating they were suppressed by the shock relative to low-firing cells.

### Local field potential (Fig. 5, 6, Supplementary Fig. S7)

LFP data on all electrodes, low pass filtered with a cut-off frequency of 2 kHz and sampled at 8 kHz during acquisition, was low pass filtered with a cut-off frequency of 200 Hz and downsampled at 500 Hz off-line for further analysis (except in Figure 8, where LFP data for ripple detection was low pass filtered at 400 Hz and downsampled at 1 kHz). For analysis of theta phase, LFP data from all electrodes was filtered between 6 and 10 Hz using the eegfilt function from the EEGLAB toolkit ^55^, first filtering using a low-cut set to 10 Hz, then a high-cut set to 6 Hz. Data was then restricted to running periods. Mean theta power was calculated on each electrode by first calculating the Hilbert transform of the data, then calculating the absolute of the resulting analytic signal. For each hemisphere, the electrode with the greatest theta power was then selected, and theta phase was calculated from the analytic signal using the *angle* function in Matlab. Spikes were assigned a theta phase using a custom circular interpolation algorithm. Spike phases were then separated into 20 equally spaced phase bins and spike probability in a bin was calculated as the number of spike phases in that bin divided by the total number of spike phases. For display purposes only, the 95% confidence intervals were constructed for the spike-phase probability plots using boostrapping with 2000 resamples with replacement.

To calculate phase locking statistics we used the CircStat Matlab toolbox ^56^. Specifically, to calculate the mean phase angle, we used the circ_mean function. To calculate the statistical significance of the differences in these two measures between two populations, we combined the spike phases from the two populations (Supplementary Fig. 6c, d). From this combined population, using random sampling with replacement, we then created two surrogate populations with the same lengths as the original populations, and calculated the differences in the mean phase angles between these two populations. This was repeated 2000 times, and we calculated 95% and 99.9% confidence intervals from the resulting null-distributions. The difference between two populations was considered significant if it fell outside the 95% confidence intervals of the null distribution.

To calculate the ratio of spikes fired on the descending wave vs. the ascending wave/peak of theta, (Fig. 5d), we divided the number of spikes with phases from 36° to 108° by the number with phases from 288° to 360° or 0° to 18°.

To calculate the phase amplitude modulation index (Figure 6a), which measures the extent to which LFP phase at a given frequency modulates power at some other frequency, power and phase on each tetrode which recorded place cells was calculated using the continuous wavelet transform with Complex Morlet wavelets (bandwidth parameter of 1.5 and a center frequency of 1) with scales corresponding from 2 to 200 Hz at 2 Hz intervals. The modulation index was then calculated at each phase frequency and power frequency as previously described ^57^. Briefly, phase at a given frequency was binned into 18 intervals. Then for a given phase bin, the mean wavelet power at each frequency was calculated. The mean wavelet power at a given phase bin was then normalized by the sum of wavelet powers across all phase bins at that phase frequency. Shannon entropy for a pair of phase and power frequencies was then calculated as

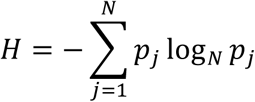

where *N* is the number of phase bins, and *p_j_* equals the normalized wavelet power at phase bin *j*. Finally, the modulation index was given by:

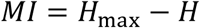

where *H*_max_ is 1.

To calculate gamma amplitude-theta phase modulation (Figure 6b), we calculated power at as above at frequencies from 20 to 200 Hz at 2 Hz intervals. Theta phase was calculated as above, and binned into 50 bins, and the power time series at each frequency was separated into the appropriate theta phase bin. In the plot, the mean power at each frequency was subtracted.

For the slow-gamma analysis (Figure 6c-h), we calculated mean power between 30-50 Hz using the continuous wavelet transform as above, excluding shock times, since we observed broad increases in power during the shock due to the shock artifact. Slow-gamma high power episodes were defined as periods of at least 20 ms duration when power was above the median power, and low power episodes were defined as periods below the median power. Spikes were separated into the two classes of slow-gamma episodes based on gamma power on the electrode on which they were recorded.

Because high power episodes clustered around earlier phases of theta, as expected, and low power episodes around later phases, the distributions had to be normalized by the distribution of theta phases during these episodes to determine the mean phases of the spike populations. For the spike distributions shown in Fig. 6c and g, this meant simply dividing the distributions by the similarly binned distributions of the theta phases. For the bootstrap analysis whose results are shown in Fig. 6d-f and h-j, for each iteration of the bootstrap, before combining the two populations of spikes, we first downsampled each population to flatten the theta phase distribution. To do this, we assigned spikes to phase bins, and then for each bin we downsampled the spikes based on the theta phase distribution for that bin. For example, if the number of theta phases at that bin was 1.5 times greater than the minimum, we randomly sampled ∼66.6% of the spikes at that bin (without replacement). The procedure then continued with these 2 downsampled populations as in the bootstrap analysis presented in Fig. 5.

To calculate the strength of phase locking for each population, we computed the mean resultant length (Fig. 6f and j; Fig. S7c and f) using the circ_r function from the CircStat Matlab toolbox, with statistics calculated as for mean phase.

### Replay analysis (Figure 7, S7)

We defined candidate awake replay events as population events with durations of at least 100 ms occurring when the animal’s velocity was less than 5 cm/sec and characterized by a peak elevation of the multi-unit firing rate of at least 3 standard deviations above the mean and a minimum firing rate of no less than half the mean. To calculate the ripple-centred firing rate (Fig. 7a, b), for each cell and population event, we calculated the mean firing rate in a 100 ms window centred on the time when peak power between 150Hz-300 Hz occurs. Each cell’s firing rate was then averaged across all population events.

The patterns of spiking during population events were decoded using a Bayesian decoding algorithm ^36, 58^. Because the choiceArm paths overlapped before and after the central choice arm, and because the spatial representation remapped across learning, we ran the decoding algorithm on each event six times using each of six templates: 1) beforeChoice-choiceArm1 pre-learning, 2) beforeChoice-choiceArm2 pre-learning, 3) beforeChoice-choiceArm3 pre-learning, 4) beforeChoice-choiceArm1 post-learning, 5) beforeChoice-choiceArm2 post-learning, 6) beforeChoice-choiceArm3 post-learning. The decoded event with the highest sequence score (see below) was selected for subsequent analysis. Each event was subdivided into overlapping 20ms windows (5ms step size). The probability of spiking activity in a given window of the event corresponding to a position on the track was given by:

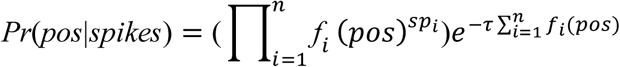

where *f_i_*(*pos*) is the value of firing rate by position vector of the *i*th unit at position *pos* in the template, *sp_i_* is the number of spikes fired by the *i*th unit, τ is the time window duration (20 ms), and *n* is the total number of cells. Posterior probabilities were normalized:

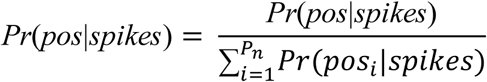

where *P_n_* is the total number of positions. Putative events were discarded if fewer than 4 place cells fired spikes. To verify that the algorithm decoded position accurately, we decoded position from spikes fired in 100 ms windows while the animal ran the task (Figure 7A). The decoded position was taken as the spatial bin with highest posterior probability in a given time window.

To determine the quality of a given replay event, we calculated a replay sequence score according to the methods of Grosmark and Buzsaki ^24^. First, the weighted mean was calculated:

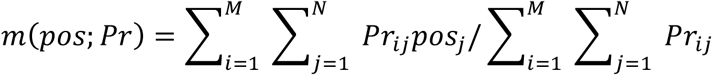

The weighted covariance was calculated:

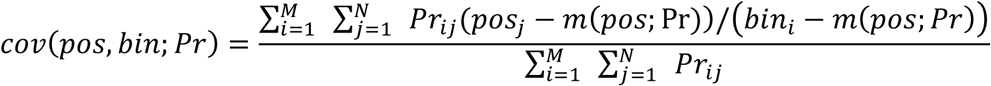

And the weighted correlation was calculated:

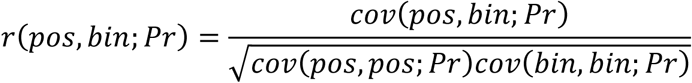

*Pos_j_* is the *j*th spatial bin, *bin_i_* is the *i*th time bin in the event, *Pr_ij_* is the Bayesian posterior probability for p*os_j_* and *bin_i_*, *M* is the total number of time bins and *N* is the total number of spatial bins.

We then decoded the events using 1000 templates in which we circularly translated unsmoothed rate maps by a random number of bins separately for each unit and then smoothed them (place cell shuffle), and 1000 template templates in which we circularly translated the population vectors at each spatial bin separately for each population vector (population vector shuffle). A weighted correlation was then calculated for each shuffled event. Because we used six template maps for decoding, we applied a correction for statistical significance testing; an event was considered statistically significant if it’s absolute weighted correlation was greater than 99.2% of absolute place cell shuffled event weighted correlations (i.e. *p* = 0.05/6 = 0.0083) and 99.2% of the population vector shuffled event weighted correlations (*p* = 0.0083).

To determine the per cell contribution (PCC), we first calculated sequence scores for each significant events as:

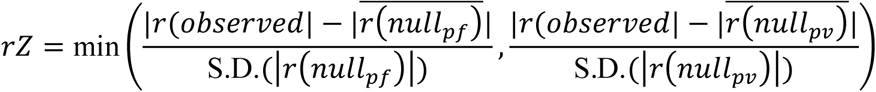

where *r(null_pf_)* and *r(null_pv_)* denote the place field and population shuffled distribution of weighted correlations, respectively. Then, for each event, the contribution of each participating cell was determined by calculating the weighted correlation for that event using a template in which only the firing rate by position vector of that unit was taken from the shuffled rate maps; if *rZ* = *rZ_pf_*, then the place field shuffled maps were used, otherwise the population vector shuffled were used. This was repeated 1000 (i.e. using a different shuffled vector on each iteration) and a sequence score corresponding to the shuffled unit and the type of shuffle (i.e. place field or population vector) as:

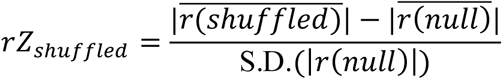

and then:

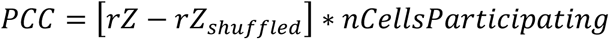

For each cell, PCC scores were averaged across all events within which it participated. In a separate analysis, a cell’s likely of participating in replay events was determined by dividing the number of significant events in which the cell fired at least 1 spike by the total number of significant events.

### Statistics

All statistical analyses were performed in Matlab, with the exception of the Friedman Test (non-parametric repeated measures ANOVA) and associated post-hoc Wilcoxon signed rank test (Figure S8) which were performed in R. Paired *t*-tests were used to assess the behavioral data in Figure 1; elsewhere, except where noted, the Wilcoxon rank sum test was used to assess the significance of difference between 2 groups, whereas the Kruskal Wallis test followed by post-hoc Dunn-Sidak tests was used for comparisons between more than 2 groups. Summary data was presented as box plots; the bottom and top of the central boxes represent the 25^th^ and 75^th^ percentiles, with the central line representing the median, whiskers extending a maximum of 1.5 * length of the 25^th^ to 75^th^ percentile distance, and any additional data points plotted as outliers.

---

**Supplementary Figure 1.**
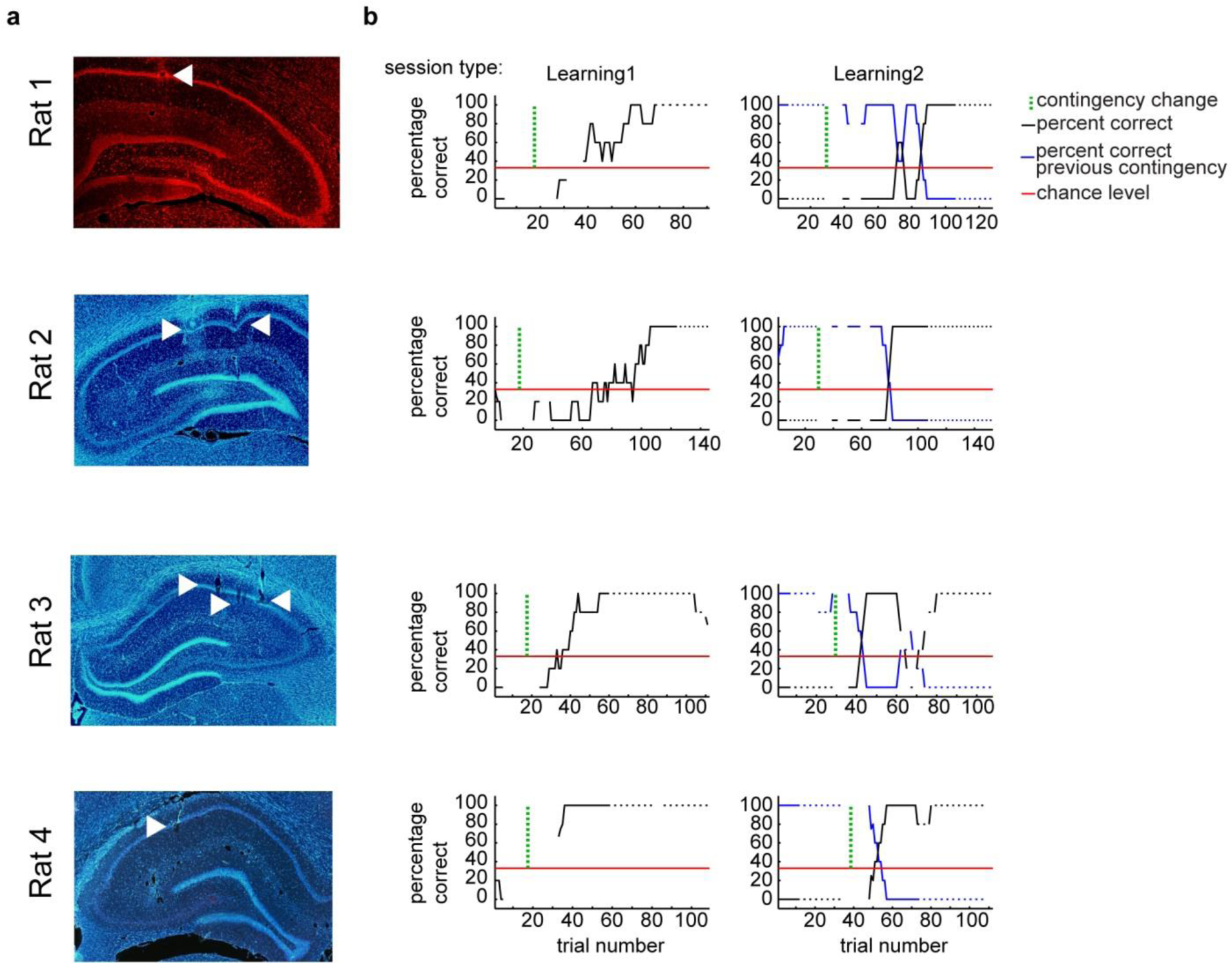
Histology and behavor. a) Representative brain slices cut from each of the four animals stained with NeuroTrace 530/615 fluorescent Nissl stain (red; Invitrogen) or DAPI (blue), showing recording locations in dorsal CA1. b) Percentage correct for each trial was calculated by averaging the five surrounding trials (i.e. current trial plus 2 immediately before and 2 immediately after). Breaks in the curve indicate forced rather than free choice trials. The “percentage correct previous contingency” in the Learning2 sessions refers to the animal’s performance in running the safe arm from the pre-contingency change epoch (also, same contingency as in the previous shockCtrl session and the post-contingency change epoch from the preceding Learning1session).

**Supplementary Figure 2.**
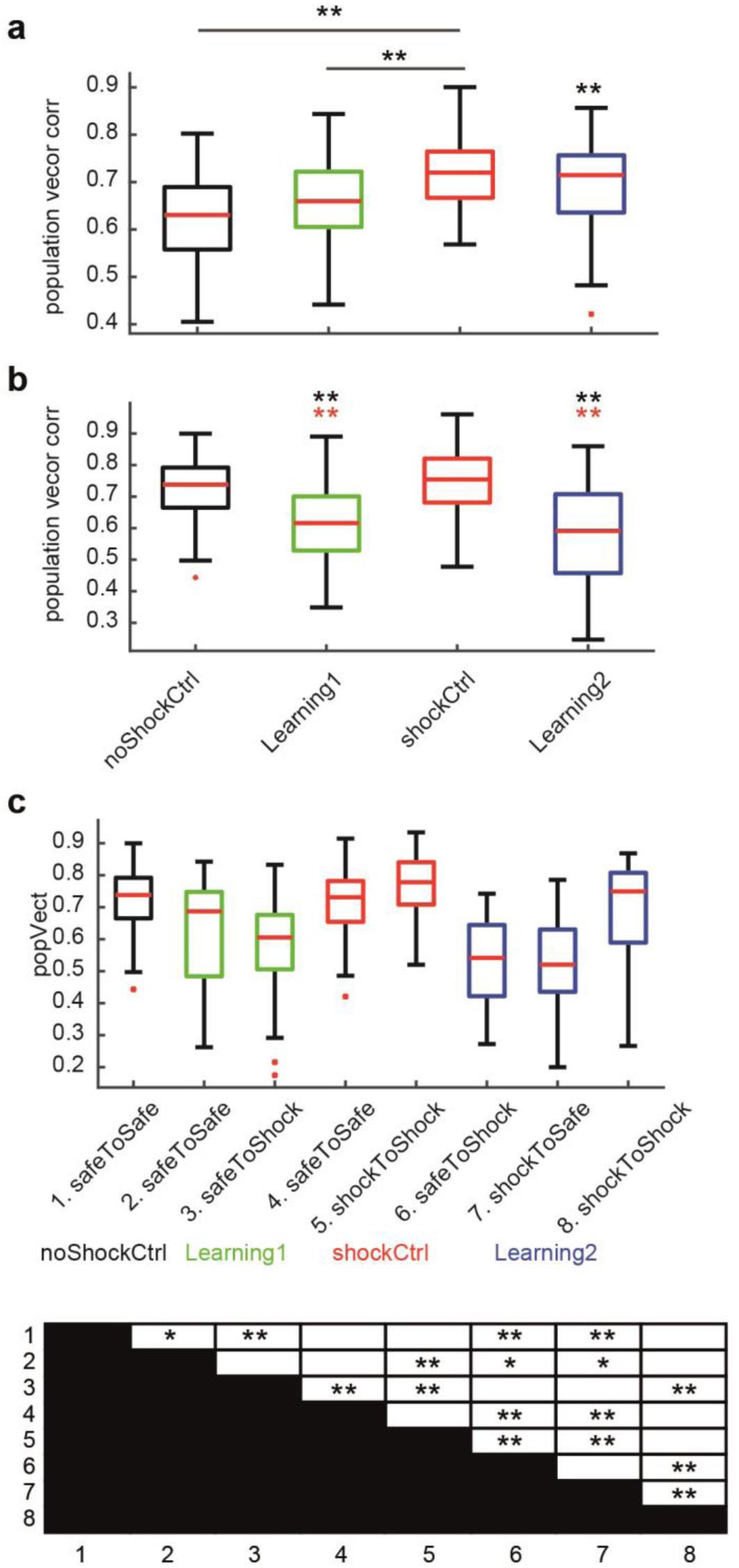
Arm specific remapping. (**a**) Boxplots showing the PV correlations between the pre- and post-rate maps on the beforeChoice path (see Fig. 1). Kruskal Wallis test, *H*(3) = 36.6, *p* < 0.001. (**b**) PV correlations between pre- and post-rate maps on the choice arms. Kruskal Wallis test, *H*(3) = 165.2, *p* < 0.001. (**c**) PV correlations between pre- and post-rate maps on each type of choice arm from each session type (top, see Fig. 1f). Kruskal Wallis test, *H*(7) = 206.1, *p* < 0.001. Bottom, summary of post-hoc Dunn-Sidak test. * and ** denote significance levels p < 0.05 and p < 0.001 respectively. ** above horizontal bar indicates significant difference between the two groups. Colored ** directly above box indicates significant difference from the group of the same color.

**Supplementary Figure 3.**
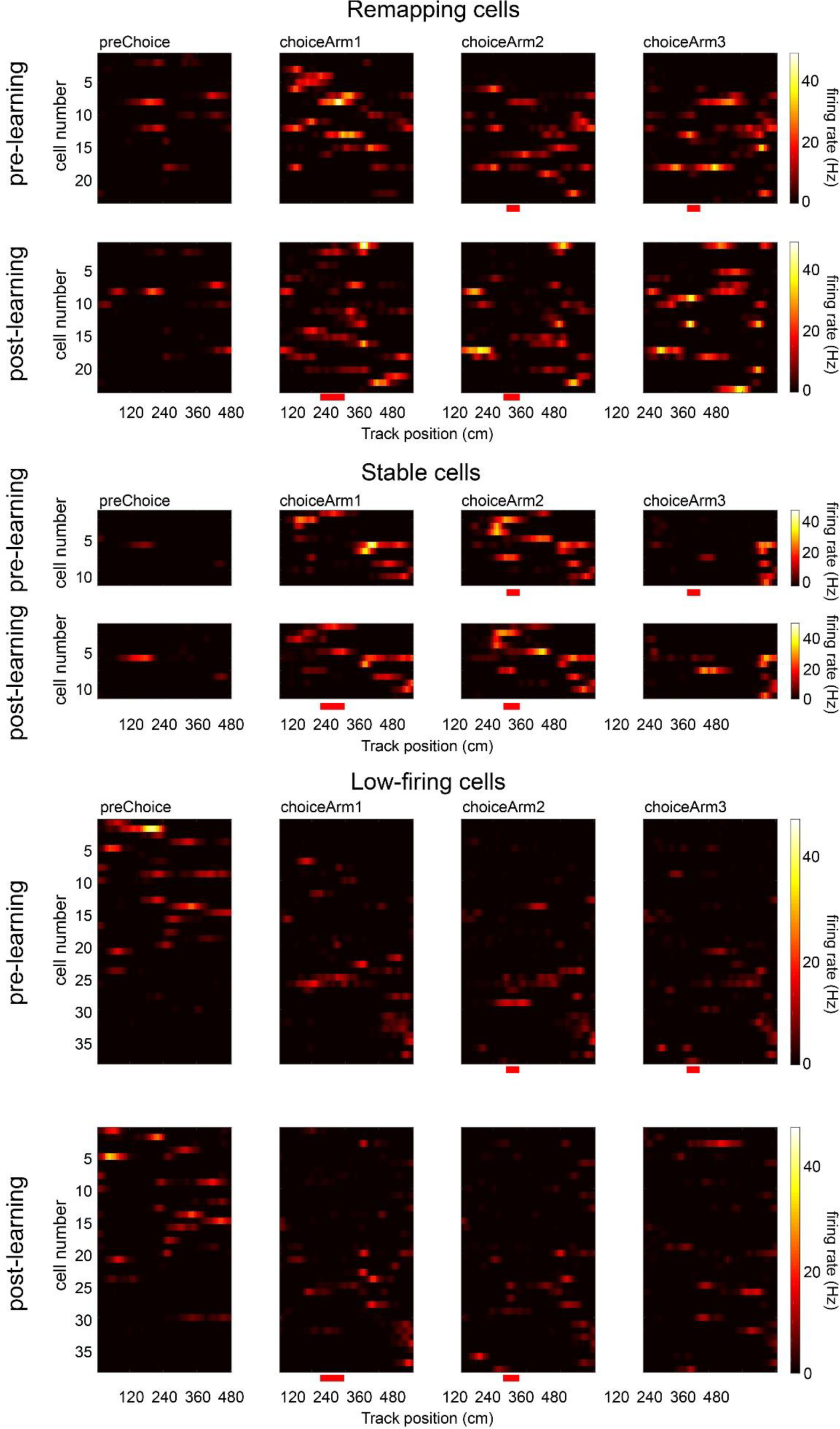
All place fields recorded in a single session. Place cells were recorded from Rat 4, shockToShock session. Fields are arranged by the center of mass of their firing distribution across the 4 combined and linearized paths (preChoice and choiceArms 1-3).

**Supplementary Figure 4.**
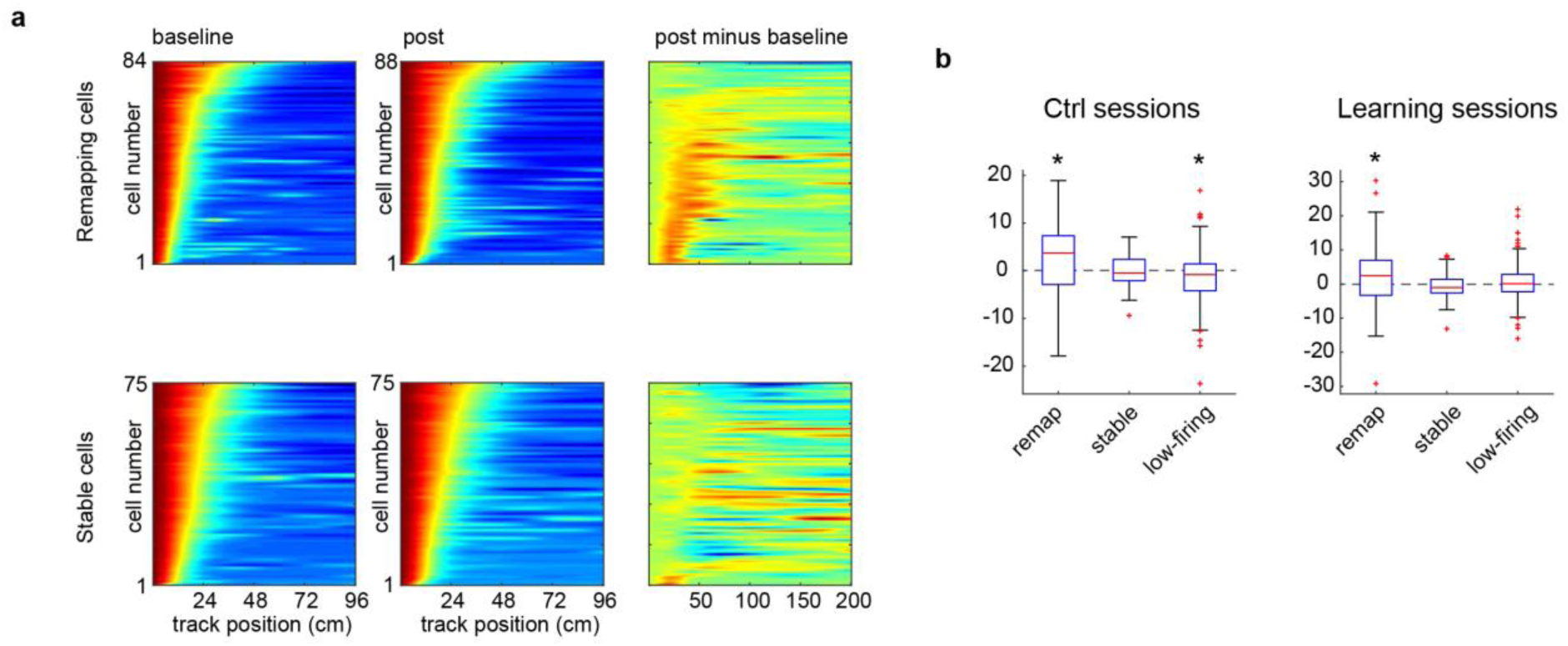
Expansion of place fields in remapping cells. (**a**) Remapping (top) and stable (bottom) cell rate map autocorrelations from learning sessions (left, baseline epoch; middle, post-learning epoch) plotted as heat maps and arranged vertically according the number of high correlation bins. Right, the baseline heat map is subtracted from the post-learning epoch, revealing that the area of high autocorrelation is larger during post-learning than during the baseline in remapping, but not stable, cells. (**b**) Difference in spatial scale from baseline to post-learning epoch in control (left) and learning (right) sessions. Wilcoxon signed-rank test, ctrl sessions: remapping, *Z* =2.09, *p* = 0.037; stable, *Z* = −0.319, *p* = 0.754; low-firing, *Z* = −2.84, *p* = 0.0045; learning sessions: remapping, *Z* = 2.07, *p* = 0.039; stable, *Z* = −1.91, *p* = 0.057; low-firing, *Z* = 0.950, *p* = 0.3419).

**Supplementary Figure 5.**
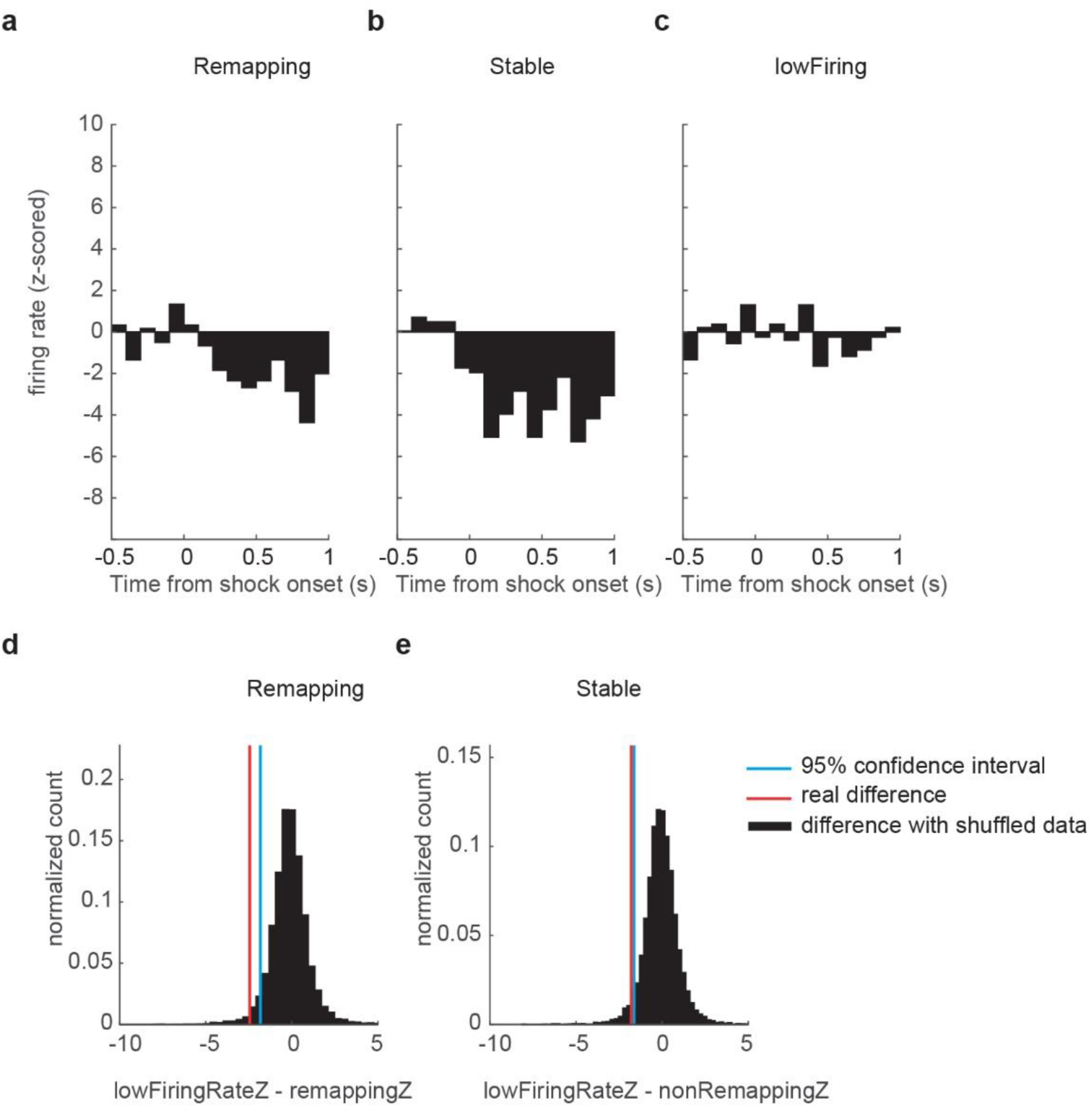
Shock-induced modulation of remapping and stable cells. (**a-c**) Z-scored peri-event time histograms (PETH) centered around shock (time 0) for (**a**) remapping, (**b**) stable, and (**c**) low-firing cells. (**d, e**) Significance of difference between (**d**) remapped and low-firing and and (**e**) stable and low-firing cells calculated using bootstrap analysis. In both cases, the differences lie outside the 95% confidence interval for the control distributions, and so are deemed to be statistically significant.

**Supplementary Figure 6.**
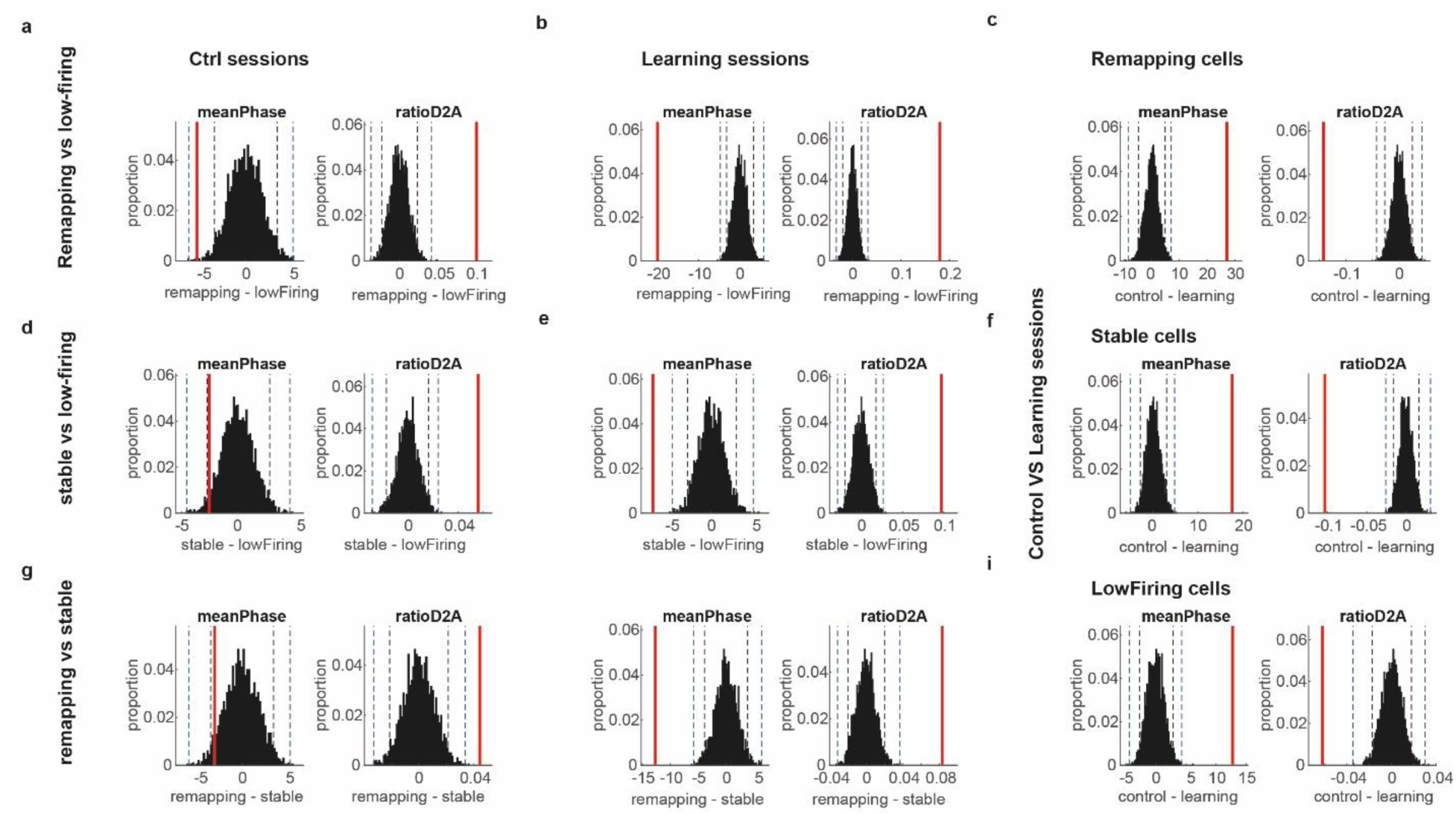
Statistical tests of theta phase preferences. (**a-i**) Differences (red) in mean phase (left) and ratio of spikes on the descending to ascending wave of theta (right) compared to control bootstrap distributions (black; 95% and 99.9% confidence intervals are denoted by dashed lines) between (a) remapping and low-firing cells during ctrl sessions, (b) remapping and low-firing cells during learning sessions, (c) remapping during control and during learning, (d) stable difference lies outside the 95% confidence intervals of the control distribution. and low-firing cells during ctrl sessions, (e) stable and low-firing cells during learning sessions, (f) stable during control and during learning, (g) remapping and stable cells during ctrl sessions, and (h) remapping and stable cells during learning sessions, (i) low-firing during control and during learning. Differences are considered statistically significant if they lie outside the 95% confidence interval.

**Supplementary Figure 7.**
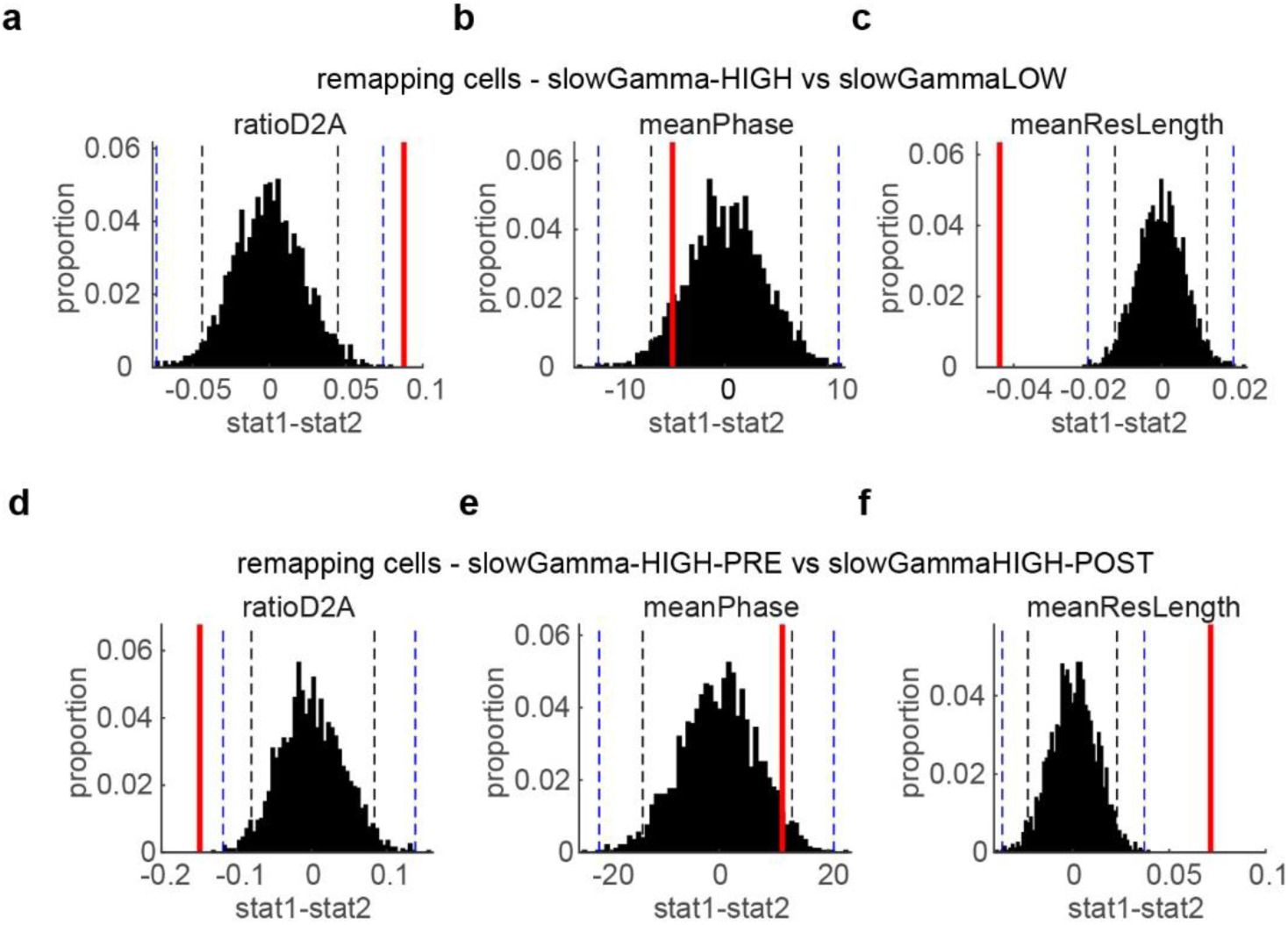
Statistical tests of theta phase modulation of remapping cell spikes by slow gamma power during Learning sessions. (**a-c**) Differences (red) in ratio of spikes on the descending to ascending wave of theta (**a**), mean theta phase (**b**), and mean resultant length (**c**; mrl) between remapping cell spikes during periods of elevated or decreased slow gamma power. (**d-f**) Differences (red) in ratio of spikes on the descending to ascending wave of theta (**d**), mean theta phase (**e**), and mean resultant length (**f**; mrl) between remapping cell spikes fired prior to (pre) or after (post) the contingency change. 95% and 99.9% confidence intervals are denoted by dashed lines. Differences are considered statistically significant they lie outside the 95% confidence interval.

**Supplementary Figure 8.**
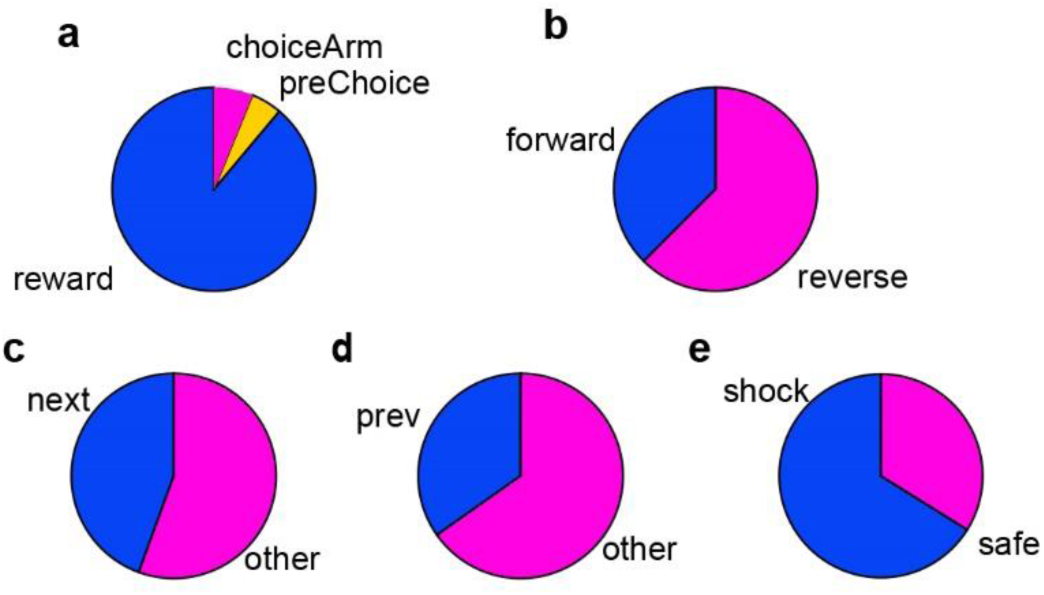
Replay event properties. (**a**) Proportion of events (*n* = x) occurring on preChoice or choiceArm paths or at either of the two reward locations (smallRwd and reward). (**b**) Proportion of forward and reverse events. Binomial test, *p* = 0.033. (**c**) Proportion of forward replay events that encoded the next path run by the animal. Binomial test, *p* = 0.49. (**d**) Proportion of reverse replay events that encoded the previous path run by the animal. Binomial test, *p* = 0.88. (**e**) Proportion of replay events that encoded shock or safe arms. Binomial test, *p* = 1.

**Supplementary Figure 9.**
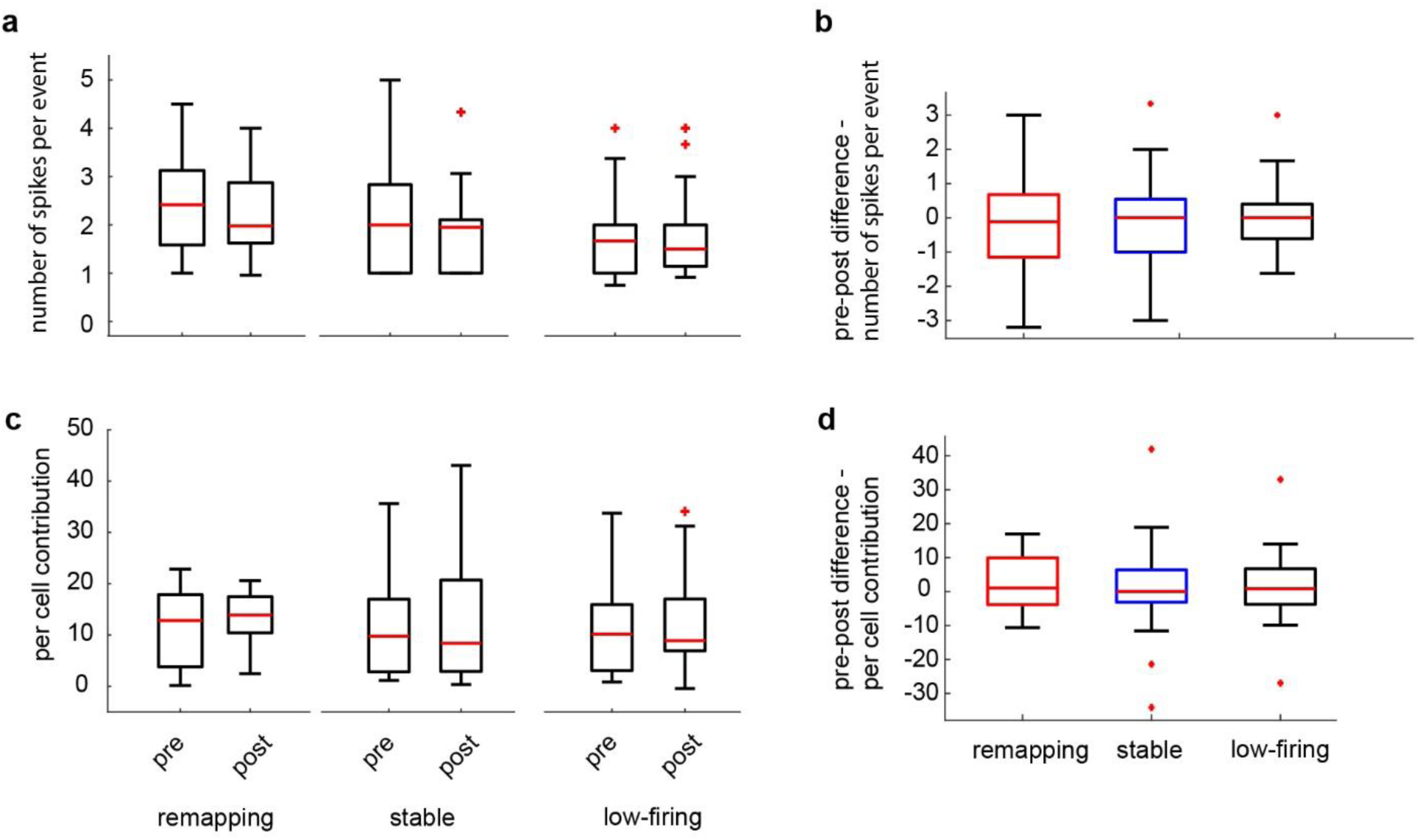
Awake replay of remapping place cells, continued. (**a**) Number of spikes fired per replay event (includes only events in which a given cell fired at least 1 spike) during pre-contingency change and post-learning epochs (Wilcoxon signed-rank test, remapping: *Z* = 0.724, *p* = 0.469; stable: *Z* = 0.577, *p* = 0.577; low-firing: *Z* = 0.252, *p* = 0.801). (**b**) Change in number of spikes fired from pre-contingency change to post-learning epochs (Kruskal Wallis test, *H*(2) = 0.323, *p* = 0.851). (**c**) Per cell contribution scores (includes only events in which a given cell fired at least 1 spike) during pre-contingency change and post-learning epochs (Wilcoxon signed-rank test, remapping: *Z* = −1.02, *p* = 0.309; stable: *Z* = −0.639, *p* = 0.523; low-firing: *Z* = −0.874, *p* = 0.382). (**d**) Change in per cell contribution scores from pre-contingency change to post-learning epochs (Kruskal Wallis test, *H*(2) = 0.215, *p* = 0.898).

